# Propagating cortical waves coordinate sensory encoding and memory retrieval in the human brain

**DOI:** 10.1101/2024.06.24.600438

**Authors:** Yifan Yang, David A. Leopold, Jeff H. Duyn, Xiao Liu

**Affiliations:** Department of Biomedical Engineering, The Pennsylvania State University, University Park, PA, 16802, USA; Neurophysiology Imaging Facility, National Institute of Mental Health, National Institute of Neurological. Disorders and Stroke, and National Eye Institute, National Institutes of Health, Bethesda, MD, 20892, USA; Section on Cognitive Neurophysiology and Imaging, Laboratory of Neuropsychology, National Institute of Mental Health, National Institutes of Health, Bethesda, MD, 20892, USA; Advanced MRI Section, Laboratory of Functional and Molecular Imaging, National Institute of Neurological Disorders and Stroke, National Institutes of Health, Bethesda, MD 20892, USA; Institute for Computational and Data Sciences, The Pennsylvania State University, University Park, PA, 16802, USA

**Keywords:** infra-slow brain waves, semantic coding, memory encoding, memory recall, global signal, brain states, hippocampal ripples

## Abstract

Complex behavior entails a balance between taking in sensory information from the environment and utilizing previously learned internal information. Experiments in behaving mice have demonstrated that the brain continually alternates between outward and inward modes of cognition, switching its mode of operation every few seconds. Further, each state transition is marked by a stereotyped cascade of neuronal spiking that pervades most forebrain structures. Here we analyzed large fMRI datasets to demonstrate that a similar switching mechanism governs the operation of the human brain. We found that human brain activity was punctuated every several seconds by coherent, propagating waves emerging in the exteroceptive sensorimotor regions and terminating in the interoceptive default mode network. As in the mouse, the issuance of such events coincided with fluctuations in pupil size, indicating a tight relationship with arousal fluctuations, and this phenomenon occurred across behavioral states. Strikingly, concurrent measurement of human performance in a visual memory task indicated that each cycle of propagating fMRI waves sequentially promoted the encoding of semantic information and self-directed retrieval of memories. Together, these findings indicate that human cognitive performance is governed by autonomous switching between exteroceptive and interoceptive states. This apparently conserved feature of mammalian brain physiology bears directly on the integration of sensory and mnemonic information during everyday behavior.

## Introduction

The human brain undergoes slow, spontaneous fMRI fluctuations during rest, in the absence of external stimulation and task engagement (1, 2). While this activity has been used widely to characterize the functional connectivity between brain regions (2–5), its contribution to the normal operation of the brain has remained elusive. Two curious features of this activity that have drawn attention in recent years are its manifestation as discrete, quasi-periodic events and its spatiotemporal propagation across the brain (6–9). Recent work describes such propagation as moving from low-order sensory-motor (SM) regions to high-order default mode network (DMN) (7, 10). This traversing of the cortical hierarchy has been compared to cross-layer error back-propagation required for optimizing artificial neural networks (11, 12), raising the prospect that these waves may play a physiological role in learning and memory consolidation.

Analogous global brain dynamics have been observed at the single neuron level in the mouse. These dynamics are associated with arousal fluctuations and manifest as massive spiking cascades involving ∼70% of recorded neurons across the forebrain and playing out over several seconds (13). During both spontaneous activity and periods of visual stimulation, spiking cascades were coordinated in time with hippocampal sharp-wave ripples (SPW-Rs), a neurophysiological event known to be involved in memory functions (14). In the case of visual stimulation, each cascade cycle involved transitioning from a phase of high-efficiency sensory encoding to a phase of heightened SPW-Rs (15). Together, these observations suggest a mechanism by which the mouse brain routinely switches between exteroceptive and interoceptive modes.

One attractive possibility is that the fMRI waves in the human brain and spiking cascades in the mouse brain reflect the same or homologous underlying neurophysiological processes. Indeed, they share common features. For example, both phenomena are manifest as quasi-period events that transpire over seconds time scales, affect global forebrain activity, and are demonstrably coupled to arousal fluctuations (13, 16). In the absence of external stimulation, fMRI waves in humans propagate between two sets of brain networks showing opposite responses to cognitive tasks (17–19), and the spiking cascade sequence in mice involves the interplay between two groups of neurons with opposite activity modulations during locomotion (13). When hippocampal SPW-Rs were measured together with concurrent fMRI in the monkey, they were synchronized with fMRI changes across the brain (20, 21). Interestingly, this mapping revealed that sensory/motor areas exhibited distinct delays from higher-order regions, suggestive cross-hierarchy propagation (20, 21). Nevertheless, it remains unknown whether propagating fMRI events are the macroscopic counterpart of neural firing cascades. More importantly, it is also unclear whether fMRI waves, like neural firing cascades, persist during stimulation or play a role in coordinating sensory and memory cycles during wakefulness.

In the present study, we analyzed multiple human fMRI and mouse neuronal recording datasets to address this topic. Similar to the spiking cascades in the mouse, the propagating fMRI waves in the human brain persisted during the performance of a visual memory task. The fMRI wave cycle alternately increased the encoding of sensory information and the efficiency of memory retrieval function across each cycle. The cascades and fMRI waves were similarly synchronized to pupil dilations in humans and mice, suggesting a shared neuromodulatory basis. These findings thus demonstrate similar, internally generated physiological cycles coordinating exteroceptive and interoceptive cognitive activity in the human and mouse brain, suggesting an evolutionarily conserved mechanism governing mammalian forebrain function.

## Results

### Fluctuating arousal entrains brain-wide events across the mouse and human forebrain

Pupil diameter is a surrogate signal for fluctuating arousal that is readily measured in both human and mouse subjects during rest (23, 24). We observed that the dynamic changes in pupil diameter were matched to the occurrence of brain-wide events in both species, thus providing a means to compare spiking cascades and fMRI waves.

In the mouse, pupil size fluctuations, indicative of changes in arousal state, were prominent during periods of immobility, with or without visual stimulation, as evident in data from the Allen Institute Visual Coding project (25) (**Fig. 1A** and **1B**). Across the brain, we found that pupil dilations coincide with moments of widespread spiking events, in which neurons fire sequentially in reproducible patterns (**Fig. 1B, C**, red arrows). The same dynamics were derived previously without reference to pupil data and described as brain-wide spiking cascades (13). We repeated the same analysis on a two-photon calcium imaging dataset (26) and another large-scale Neuropixel dataset with broader coverage of mouse brain (27), revealing that these pupil-associated cascades span across widespread brain regions and involve multiple neuron subtypes (**Fig. S1** and **S2**).

**Figure 1.**
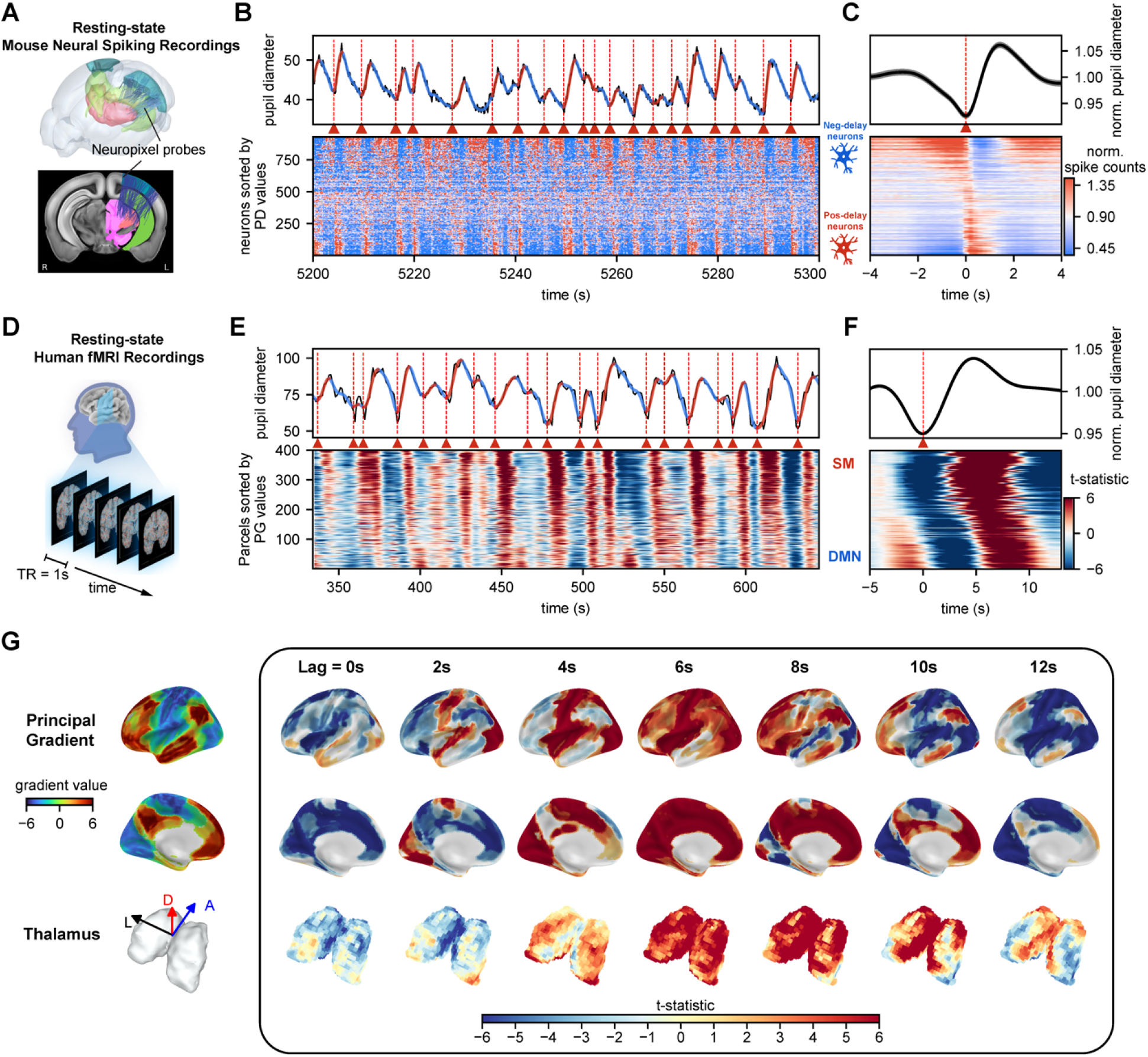
Association of spontaneous pupil dilations and brain-wide sequential activity in the mouse and human brain. (A) Locations of neuropixel probes from all mice in the Allen Mouse Brain Common Coordinate Framework with major recording sites are color-coded: visual cortex (blue), hippocampus (green), and thalamus (pink). Top: Three-dimensional illustration of the probe insertion in mouse brain. Bottom: Two-dimensional projection of the probes onto a middle brain slice. (B) Close coordination of spontaneous pupil dilation and spiking cascade occurrence in the mouse brain during 100 s of stationary visual stimulation. Top: spontaneous fluctuation of pupil diameter with alternating dilation (red) and constriction (blue) phases, with the onset of dilation marked by red dashed lines and triangular symbols. Bottom: normalized spiking activity of all recorded neurons that are sorted according to the principal delay profile, revealing the correspondence between single pupil dilations and spiking cascades of sequential activations from negative-delay neurons (blue symbolic neurons) to positive-delay neurons (red symbolic neurons). (C) The representative mouse’s normalized pupil diameter (top) and neuronal spiking activities (bottom) averaged around the onset of pupil dilation over an 8 s window. (D) Schematic of a resting-state fMRI scan from the Human Connectome Project 7-Tesla (HCP-7T) dataset. (E) Close coordination of spontaneous pupil dilation and propagating fMRI waves in the human brain during 310 s period of rest, similar in layout to (B). Pupil diameter fluctuation (top) and concurrent fMRI signals of various brain regions are sorted by their principal gradient value (22) (bottom). (F) The averaged pupil diameter (top) and fMRI signals (bottom) at the onset of pupil dilation over an 18-sec time window, summarized from all 184 subjects. (G) The pupil-dilation-associated fMRI changes mapped onto the brain’s surface. The maps were shown for 7 evenly spaced time lags, from 0 to 12 seconds following the onset of pupil dilation. The first and second rows display maps of the brain’s cortical surface, and the third row presents the thalamic volume map. Directional arrows denote dorsal (D), anterior (A), and anatomical left (L) directions.

In the Human Connectome Project (HCP) 7T dataset (28), we similarly found that pupil size changes correlated with spontaneous resting fMRI fluctuations across the brain (**Fig. 1D** and **1E**). Alignment to pupil dilation onset revealed a temporal sequence of fMRI changes progressing along a principal gradient (PG) direction (**Fig. 1F** and **S3**), which approximates the cortical hierarchy gradient (22). These events were manifest as infra-slow (multi-second) waves moving gradually from SM to DMN regions. The cortical changes were accompanied by corresponding thalamic changes (**Fig. 1G and S3F**). Such SM-to-DMN propagating waves have been identified previously without pupil data (7, 10, 22). Thus, spontaneous pupil dilations during immobile rest are associated with sequential brain dynamics of global involvement, observed as spiking cascades in mice and propagating fMRI waves in humans. The correspondence between the mouse cascade and human fMRI waves is further supported by similar changes in delta-band (1-4 Hz) activity across their cycles (**Fig. S4**).

While the function of these brain-wide events is poorly understood, evidence in the mouse ties spiking cascades to alternating periods of stimulus coding and memory operation (15). Might the fMRI waves in humans similarly regulate this switch between exteroceptive and interoceptive modes of brain function? To address this question, we investigated the occurrence of spontaneous propagating fMRI waves as human subjects performed a cognitive task involving memory. Specifically, we asked whether the sensory encoding of stimuli and successful memory retrieval performance varied as a function of these spontaneous events.

### Visual stimulus encoding predicts subsequent memory function

In order to systematically investigate the role of propagating SM-to-DMN waves on human cognition, we first needed to establish a reliable means to evaluate the encoding of visual stimuli from fMRI responses across the brain. We developed a method to do this using the Natural Scenes Dataset (NSD) (30), in which a series of 10,000 captioned natural images were shown, in the form of 4-s trials, to each of 8 subjects with each image being presented three times over 40 scan sessions on different days. For each trial, the subjects needed to indicate whether they had seen the stimulus before (**Fig. 2E**).

**Figure 2.**
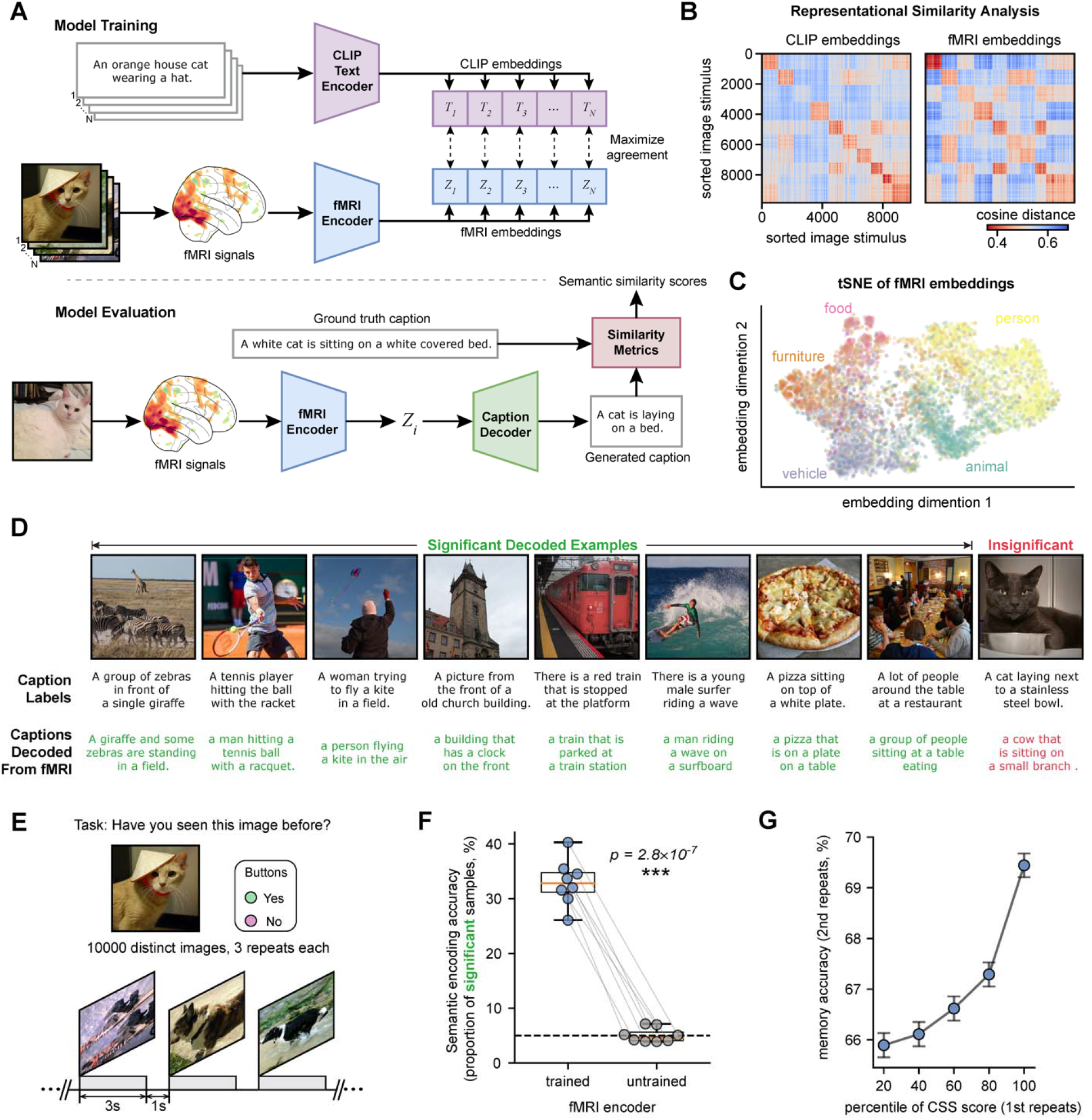
Stimulus information encoding was assessed through fMRI responses using a deep learning-based model. (A) Framework for training and evaluating the CLIP-based semantic decoder. In the training phase, an fMRI encoder is trained to map stimulus-evoked fMRI response to CLIP embedding space by maximizing the similarity between actual pairings of fMRI embedding and CLIP text embedding, while minimizing the similarity of embeddings of incorrect pairings via contrastive learning. For valuation, through the trained fMRI encoder, image-evoked fMRI responses are first encoded to fMRI embeddings in CLIP space and decoded by a pre-trained caption decoder (29) to generate text descriptions. The similarity between the generated text and ground truth text serves as an approximate but objective measure of the amount of semantic information being successfully encoded in fMRI responses, and thus of the brain’s semantic encoding accuracy. (B) The representational similarity analysis (RSA) confirms a successful training of the semantic decoder. Representational dissimilarity matrices based on cosine similarity for the semantic CLIP embeddings (left) and fMRI embeddings (right) showed highly similar structures. (C) Visualization of fMRI embeddings using t-SNE (t-distributed Stochastic Neighbor Embedding) reveals categorical distinctions within the embeddings, with each category distinctly color-coded. (D) Examples of significantly and insignificantly decoded samples, from top to bottom showing the image stimuli, the corresponding ground truth captions, and the caption decoded with evoked fMRI response. (E) Schematic of task design in the NSD dataset. Each of the 8 participants viewed 10,000 distinct images with each image randomly displayed three times across 30-40 scan sessions over a year. The stimuli presentation followed an event-related design comprising 4-second trials with 3 seconds of presentation and 1 second of baseline. (F) A box plot comparing the encoding accuracy, i.e. the proportion of significantly decoded samples based on CSS scores, between the fMRI encoder pre- and post-training. Each dot represents an individual participant, with the dotted line indicating a 5% chance level. Statistical significance is assessed using a two-sided pair-wise t-test (N=8).(G) The influence of initial image presentation encoding accuracy on subsequent memory task performance. Stimuli are binned based on the percentile of their first encoding accuracy, incremented by 20%, and the memory task accuracy of their second presentation is averaged within the bins. The results are obtained by pooling the data from all subjects.

To quantify the level of sensory stimulus encoding, we developed a novel deep learning model to decode semantic information of each image stimulus based on fMRI responses it evoked (**Fig. 2A and S5A**). The model comprised an fMRI encoder, which extracted latent representation from the fMRI responses, i.e., the fMRI embeddings, and a caption decoder (29), which translated the fMRI embeddings into descriptive text captions. The fMRI encoder was trained to align the fMRI embeddings with the contrastive language-image pre-training (CLIP) embedding space (31) through contrastive learning (32). We then quantified the semantic similarity between the fMRI-decoded caption and the original caption by a composite semantic similarity (CSS) score to measure the accuracy of semantic information encoding (see **Methods** for more detail).

Our deep learning model successfully decoded the semantic information associated with the visual stimuli based on the fMRI responses they evoked. The representation similarity analysis confirmed the alignment between fMRI and caption embeddings (**Figs. 2B, S5B**, and **S5C**), and the fMRI embeddings after training are organized as distinct categories in a low-dimensional space (**Fig. 2C**) (33). The trained model generated captions significantly similar to the ground truth captions for 33.0 ± 4.2% (mean ± SD) trials (see **Fig. 2D** and **Fig. S6A** for examples) as compared with the 5% chance-level performance of the untrained model (**Fig. 2F**, *p* = 2.8×10^−7^; and **Fig. S6B**). In the context of the cognitive task, the semantic encoding accuracy faithfully predicted subsequent memory performance: a higher CSS score at the first appearance of an image stimulus led to a higher rate of correctly recalling it at its second repeat (**Fig. 2G** and **S7**). Given this tool, it was next possible to evaluate whether the occurrence of spontaneous propagating fMRI waves might bear on the quality of stimulus encoding, subsequent memory recall, or both.

### Alternating stimulus encoding versus memory recall during propagating fMRI waves

To address the role of the propagating fMRI waves on encoding and memory performance, we first established their presence during the cognitive task. These waves were identified directly from task fMRI data without using pupil data (7) (**Fig. 3A**). Similar to the resting state, pupil diameter fluctuations remained closely tied to the occurrence of propagating waves, despite also being affected by other task events to a lesser extent (**Fig. S8A**). The duration of the SM-to-DMN waves (∼10-15 seconds) is much longer than the task trials (4 seconds), and their occurrence, propagation, and relationship to pupil fluctuations are dissociated from the structure of the concurrent cognitive task (**Fig. S8B**).

**Figure 3.**
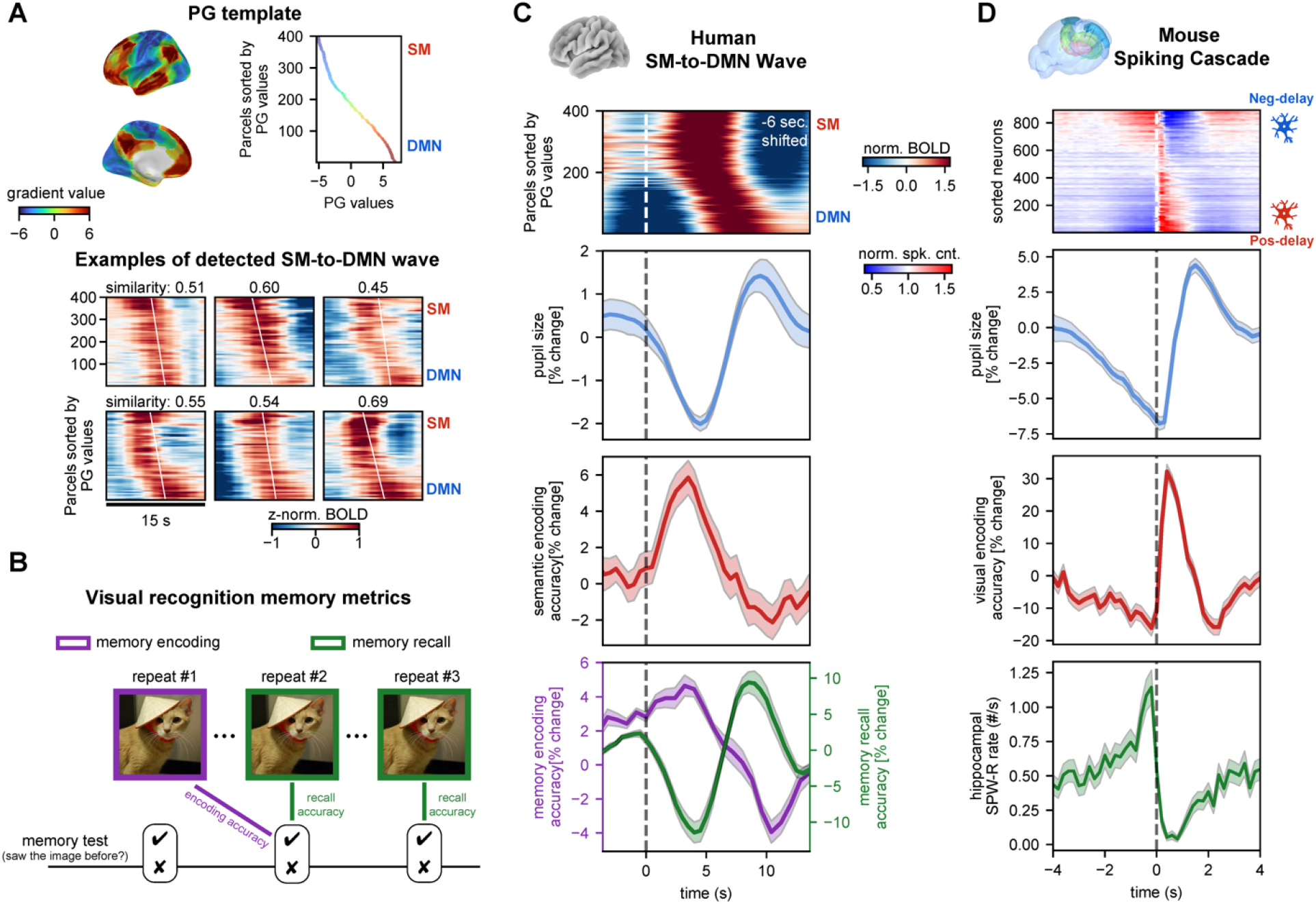
Semantic encoding and memory retrieval are oppositely modulated over the fMRI SM-to-DMN wave cycle. (A) Detection of SM-to-DMN propagating waves during the memory task. The waves were detected using template-matching methods (7). The principal gradient (PG) map was used as the template, and six examples of detected propagations are shown, with similarity values indicating the correlation between the PG template and the delay profile of fMRI segments. (B) Schematics of two distinct memory metrics. Each image has three repeats. The fMRI wave phase at the first repeat was linked to the accuracy of correctly recognizing it at the second appearance to quantify the effect of this wave dynamic on memory encoding. Then, the fMRI wave phase at the second and third repeats were linked to the accuracy of recognition tasks at the same time to quantify the effect of the wave dynamic memory recall. (C) Opposite modulations of semantic/memory encoding and memory retrieval over the cycle of the SM-to-DMN waves. The first row shows the averaged pattern of the detected SM-to-DMN waves, which was shifted backward in time by 6 seconds to account for the known hemodynamic response delay. The time zero was marked at the onset of the global mean signal increase (dashed line), which appears to correspond to the cascade center (D) as judged from the timings of the upswing in semantic and visual encoding accuracies for human and mouse data respectively. Both pupil size (second row) and memory retrieval (green in the bottom row) change significantly across the wave cycle and peak at the DMN-activated phase, whereas the semantic encoding accuracy (third row) and memory encoding accuracy (purple in the bottom row) are modulated in an opposite manner. Time series data are provided as the mean ± SEM for eight participants (N=8). (D) Opposite modulations of the visual encoding accuracy (second row) and hippocampal SPW-R rate (bottom row) across the spiking cascade cycle (first row) during stationary periods with continuous natural image stimulation in mice, adapted from (15). Time zero is marked by a dashed line indicating the onset of positive-delay neuron firing. The time series data is shown as mean ± SEM for 20 mice (N=20).

To evaluate whether encoding efficiency was influenced by propagating fMRI waves, we first used the fMRI deep learning-based decoding method described above to characterize the quality of encoding with each stimulus presentation. We found that the accuracy of such encoding varied systematically across the SM-to-DMN propagation cycle (**Fig. 3C**, red trace). Accounting for hemodynamic delays (see Methods), the stimulus encoding was strongest at the SM-activated phase of the propagating wave.

We also used memory performance as a means to assess how fMRI waves affected both stimulus encoding and memory recall. For encoding, accurate memory of individual stimuli during their second appearance was taken to indicate strong encoding at the initial presentation, whereas failure to remember a stimulus was taken to indicate weak encoding. This measure also realized the important role of the spontaneous propagating SM-to-DMN waves. Namely, the strongest memory encoding occurred when the initial stimulus (Repeat #1 in **Fig. 3B**) was presented at the SM-activated phase (**Fig. 3C**, purple trace), thus matching the fMRI deep learning-based measure of stimulus encoding described just above (**Fig. 3C**, red trace). By contrast, evaluation of recall performance, which was done at the 2^nd^ and 3^rd^ presentations of a stimulus, revealed a peak performance later in the wave cycle, when the subject recall coincided with the DMN-activated phase (**Fig 3C**, green trace). These results were similar for both short-term and long-term memory types (**Fig. S9B and S9C**).

These cyclic modulations of stimulus encoding and memory recall in humans resembled analogous observations in mice during different phases of the spiking cascades (15) (**Fig. 3D**). Specifically, the SM-activated phase of the fMRI wave matched a period within the cascade cycle (0–0.5 sec) of improved stimulus encoding, whereas the DMN-activated phase aligned with a different period within the cascade cycle (0.5–2 seconds) of increasing hippocampal SWP-R rate, which was also associated with pupil dilation (**Fig. 3C** and **3D**). While the hippocampal SWP-R rate and memory performance are clearly different measures, they may point to similar processes that transpire during more introspective modes of brain activity, commonly associated with activity of the DMN (34, 35).

### Visual semantic information coding in multiple brain regions is similarly modulated by the SM-to-DMN wave cycle

Repeating the semantic decoding using only regional fMRI data suggested that the semantic information was encoded across a wide range of brain regions, with the highest encoding accuracy observed in the visual cortex (**Fig. 4A and 4B**). Importantly, the encoding accuracy was modulated in all these regions over the SM-to-DMN wave cycle (**Fig. 4C** and **Fig. S10**) in a similar way as the whole-brain finding (**Fig. 3C**). Interestingly, the DMN, particularly its C division that encompasses the hippocampal complex and adjacent to visual association areas, exhibited the peak encoding accuracy at the SM-activated phase of the wave when its activity is not peaked, suggesting a dissociation between sensory encoding and regional activation level. These region-specific results on visual semantic encoding are consistent with those on cascade-dependent visual encoding (15), further suggesting that the spiking cascades and cross-hierarchy waves represent the same neurophysiological process conserved across mice and humans.

**Figure 4.**
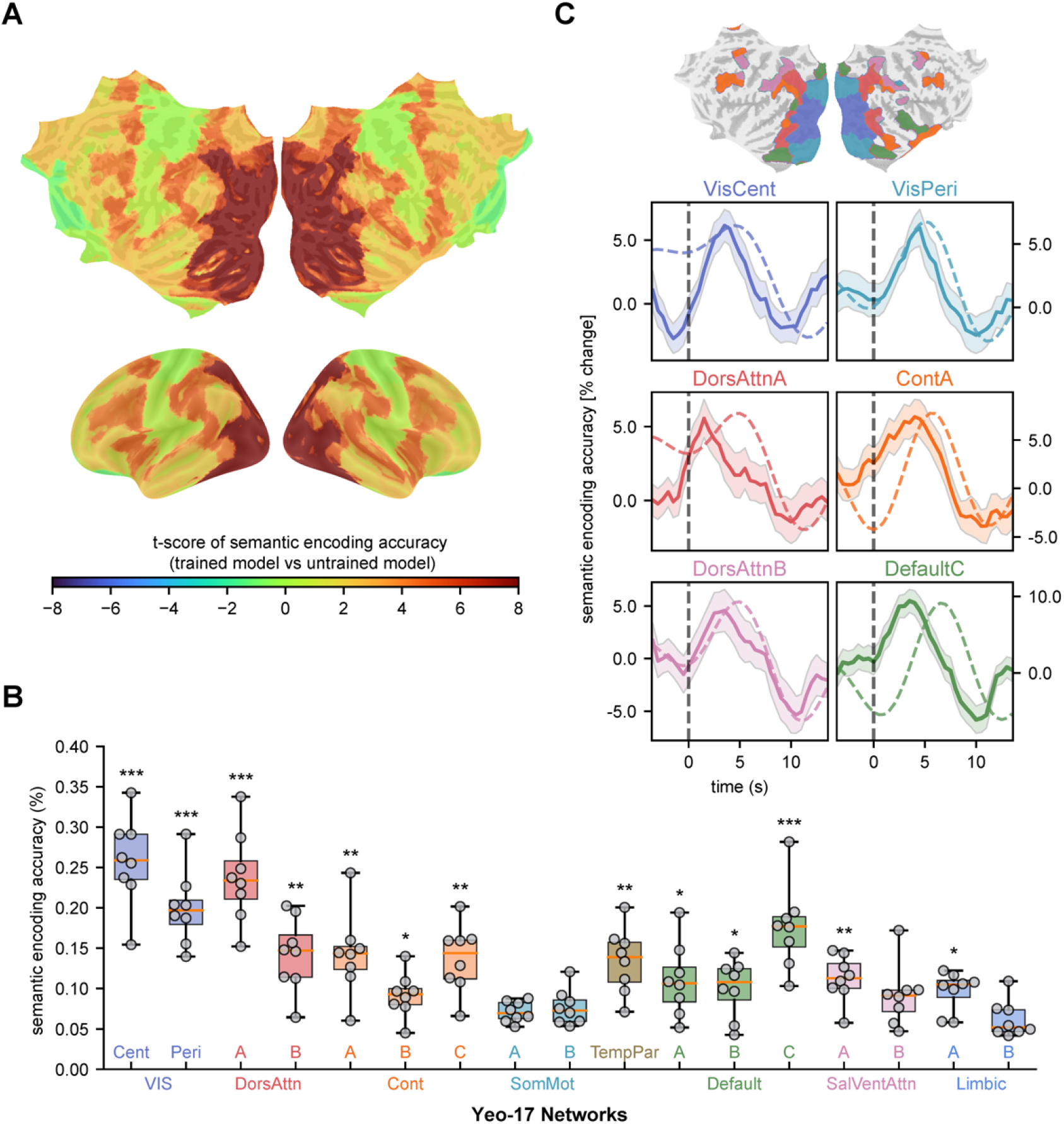
Stimulus encoding and its modulation across the SM-to-DMN wave cycle in different brain regions. (A) Cortical surface map showing the regional significance (paired t-tests: trained vs. untrained) of semantic encoding accuracy. (B) A box plot showing the semantic encoding accuracy estimated for different brain regions defined in the Yeo-17 networks atlas (36, 37). Asterisks denote the levels of statistical significance (paired t-test): *, 0.01 <p <0.05; **, 0.001 <p <0.01; ***, p <0.001. Each dot represents an individual participant. (C) Semantic decoding accuracy (solid line) is consistently modulated over the SM-to-DMN wave cycle in six brain regions exhibiting the most significant decoding accuracy. For comparison, the average activation for each region is marked with a dotted line, which is shifted ahead of time by 6 seconds to account for the hemodynamic response delay. These regions are distinctly color-coded and their locations are indicated on a flattened cortical surface. Time series data are provided as the mean + SEM for eight participants (N=8).

## Discussion

Here we showed that slow activity waves propagating over the cortical surface are associated with a counter-acting modulation of encoding and retrieval of information conferred by visual stimuli. By analyzing electrophysiological and fMRI measures of brain activity, we first demonstrated that spontaneous pupil dilations are similarly accompanied by spiking cascade dynamics in mice and SM-to-DMN propagating waves in humans, thereby unifying these two types of infra-slow (<0.1Hz) global brain activity across different spatial scales and species. Assessing the semantic encoding of visual stimuli using a CLIP-based deep learning model, we found that the SM-to-DMN propagating waves persisted during task performance and were associated with counter-valent modulation in both encoding and retrieval of the stimulus content. The encoding of semantic information and memory peaked at the early phase of SM-activation, whereas memory retrieval accuracy reached the maximum at the DMN-activated phase. Together with previous findings from mice, these results suggested that the highly structured infra-slow global brain activity serves as an evolutionarily conserved mechanism by which the brain orchestrates the execution of exteroceptive sensory sampling and internal mnemonic processes on the timescale of seconds.

The brain’s response to identical sensory stimuli is known to vary over time even on the timescale of seconds. Previous studies have shown how pre-stimulus ongoing activity and arousal state may contribute to this variability (24, 38–45). Our findings align with and extend these previous reports. Leveraging recent advances in deep learning techniques, our study goes beyond a simple quantification of response amplitude (2, 42, 43) and assesses the accuracy of the brain’s encoding of semantic information. Importantly, most previous studies have presumed that ongoing brain activity and changes in arousal occur spontaneously and randomly. As a result, much focus has been on the response modulation of ongoing activity that is temporally locked (prior to the stimulus) and spatially restricted (confined to the same local brain region). In contrast, we consider the effects of internal fluctuation in the context of highly structured brain dynamics (i.e., the spiking cascade or propagating wave) involving the large-scale coordination of activity. The initiation of these recurring global brain events is independent of visual stimulation and memory tasks, and it modulates sensory processing quasi-periodically in a continuous and persistent way.

We further found that memory retrieval was modulated over the SM-to-DMN wave cycle in a manner opposite to that of stimulus encoding, matching our previously observed counter-modulation of hippocampal SPW-R rate and visual encoding (15). This previous study did not, however, identify specific memory functions or other cognitive operations associated with SPW-Rs during the task, since SPW-Rs are usually observed during rest and sleep and often linked to offline memory consolidation (21, 46, 47). By comparing our human study results with these prior findings in mice, we found a correspondence between the cascade phase of high SPW-R rate to the wave phase of fMRI DMN activation, which is associated with a better performance in memory retrieval. This observation largely agrees with a series of recent studies on different species that linked SPW-Rs during tasks to memory retrieval (48–50), as well as the marked fMRI DMN activations (34, 35, 51, 52).

The observed modulation of sensory and memory functions over the cascade/wave cycle may be associated with a change in the direction of information flow, particularly between the cortex and hippocampus. Memory retrieval during tasks and memory consolidation during rest and sleep likely require information flow from the hippocampus to the cortex, whereas the encoding of sensory information and memory would be facilitated by a reversed flow (21, 53, 54). Thus, the cascade/wave phases optimized for sensory encoding and memory retrieval may be dominated by opposite directions of information transmission, which may rely on distinct spatial gradients in activation level. In fMRI, such activation gradients are obvious for SM-to-DMN waves with dominant SM or DMN activation at different phases. This is less clear for cascades, since the negative- and positive-delay neurons were found in all recorded brain regions (13). However, the hippocampal regions, especially CA1 and the dentate gyrus (DG), contain a much higher number of negative-delay neurons compared to any other areas, including all visual areas, whereas the thalamus has the least. Thus, the activation gradient between these two neuronal groups can be translated into spatial gradients among the hippocampus, cortex, and thalamus. We hypothesize that these gradients, alternating on the multi-second scale, determine the dominant direction of information flow, which itself occurs on much faster (millisecond) timescales. This hypothesis remains to be tested by future studies. It is worth noting that artificial neural networks also feature alternating forward/backward information flows across hierarchical layers during training (11, 12), which may thus represent a mechanism essential to the learning of all connection-based intelligence systems.

The cascade and wave dynamics reported here may represent a fundamental mechanism by which the brain coordinates the opposing operations of exteroceptive sensory sampling and internal mnemonic processes. A balance between these processes is essential for optimized cognitive performance and is likely reached under states of intermediate arousal (55, 56). Highly aroused states could break this balance by terminating this infra-slow global dynamic. Locomotion, presumably associated with heightened arousal, has been found to replace cascade dynamics with sustained firing of the positive-delay neurons (15) that are expected to promote sensory and memory encoding but impede memory retrieval (57, 58). Toward the other end of this spectrum, during drowsiness, the infra-slow global dynamic may prolong the memory consolidation phase whereas hinder encodings. The SM-to-DMN waves have been found to occur more frequently during various sleep stages and be associated with learning-related features (i.e., the rapid eye movements and possibly Ponto-Geniculo-Occipital (PGO) waves) during rapid eye movement (REM) sleep (59). Though not directly focused on the cascade and waves, recent studies convergingly point out an essential role of infra-slow neural dynamics in learning and memory. In addition to hippocampal SPW-Rs, infra-slow dynamics have been found to simultaneously coordinate the density of sleep spindles, an electrophysiological feature that has relevance for learning and memory (60, 61). Importantly, the amplitude of infra-slow dynamics during sleep, defined through spindle density and cardiac rate, is not only correlated with memory performance on the subsequent day (62), but optogenetically enhancing it also leads to improved memory (63). Similar to the modulation of brain activity during cascades and waves described here, such spindle-based infra-slow dynamic alternates between an offline phase, characterized by higher spindle and hippocampal SPW-Rs rates with low arousal, and an online phase, marked by lower spindle and ripple rates with higher arousal and susceptibility to external stimulation (62).

The SM-to-DMN propagating wave and its effect on sensory and memory functions may offer explanations for some previous task fMRI observations. Graph-theory metrics based on fMRI connectivity/correlations, such as cartography and network flexibility, have been used to quantify brain dynamics and found associations with various cognitive components, particularly learning (64–66). Most of these metrics focused on assessing the integration and segregation of the large-scale networks, which are expected to be profoundly affected by the presence of the global SM-to-DMN waves. Thus, the waves could be an important contributor to these metrics of network dynamics. Another related phenomenon is the so-called encoding/retrieval flip, in which the deactivation and activation of the posteromedial cortex, a key component of DMN, are preferentially associated with successful memory encoding and retrieval respectively (67–69). This phenomenon can be explained by our finding that memory encoding and recall were oppositely modulated over the wave cycle with distinct DMN activations. Importantly, the present study expands this early research by incorporating the previous findings into the framework of highly structured cross-hierarchy propagating waves, which persist under various brain conditions beyond tasks.

Finally, the SM-to-DMN waves may also relate to memory dysfunction in Alzheimer’s disease (AD). The global mean BOLD (gBOLD) signal, whose peaks the SM-to-DMN waves are manifested as, has been repeatedly linked to various AD pathologies (70–72). The gBOLD peaks (also SM-to-DMN waves (16)) have been found to be coupled by strong cerebrospinal fluid (CSF) movements, known to be essential for peri-vascular waste clearance (73–75). The strength of this gBOLD-CSF coupling is indeed associated with the accumulation of amyloid-beta and tau (71, 72). Particularly, the failure of the SM-to-DMN waves to reach the DMN appeared to account for preferential amyloid-beta accumulation at these higher-order regions at the early stage of AD (71). Besides the toxic protein accumulation, AD also features dysfunctions in memory and subcortical neuromodulatory systems (76–79), which are both linked to the cascades and global waves (10, 13). Thus, it is possible that changes in this infra-slow global dynamic may also related to the dysfunction of the memory and arousal systems in AD.

## Acknowledgements

This work was supported by the Brain Initiative award (1RF1MH123247-01), the NIH R01 award (1R01NS113889-01A1), and the Intramural Research Program of the National Institute of Mental Health (ZIA-MH002838).

## Data and materials availability

For mice single neuron analysis, we used the Neuropixels Visual Coding Neuropixels and two-photon calcium imaging datasets from the Allen Institute (25, 26), accessible at https://portal.brain-map.org/overview. For resting-state human fMRI analysis, we used HCP-7T dataset from https://www.humanconnectome.org. We shared our EEG-fMRI dataset at https://openneuro.org/datasets/ds003768. For task human fMRI analysis, we used NSD dataset available at https://naturalscenesdataset.org. The Python code that produced the major results of this paper will be available at https://github.com/psu-mcnl/fMRI-Arousal.

## References

1. B. Biswal, F. Zerrin Yetkin, V. M. Haughton, J. S. Hyde, Functional connectivity in the motor cortex of resting human brain using echo-planar mri. Magn Reson Med 34, 537–541 (1995).

2. M. D. Fox, M. E. Raichle, Spontaneous fluctuations in brain activity observed with functional magnetic resonance imaging. Nat Rev Neurosci 8, 700–711 (2007).

3. M. Yu, O. Sporns, A. J. Saykin, The human connectome in Alzheimer disease — relationship to biomarkers and genetics. Nat Rev Neurol 17, 545–563 (2021).

4. S. M. Smith, et al., A positive-negative mode of population covariation links brain connectivity, demographics and behavior. Nat Neurosci 18, 1565–1567 (2015).

5. D. Zhang, M. E. Raichle, Disease and the brain’s dark energy. Nat Rev Neurol 6, 15–28 (2010).

6. T. Matsui, T. Murakami, K. Ohki, Transient neuronal coactivations embedded in globally propagating waves underlie resting-state functional connectivity. Proceedings of the National Academy of Sciences 113, 6556–6561 (2016).

7. Y. Gu, et al., Brain Activity Fluctuations Propagate as Waves Traversing the Cortical Hierarchy. Cerebral Cortex 31, 3986–4005 (2021).

8. B. Yousefi, J. Shin, E. H. Schumacher, S. D. Keilholz, Quasi-periodic patterns of intrinsic brain activity in individuals and their relationship to global signal. Neuroimage 167, 297–308 (2018).

9. B. Yousefi, S. Keilholz, Propagating patterns of intrinsic activity along macroscale gradients coordinate functional connections across the whole brain. Neuroimage 231, 117827 (2021).

10. R. V Raut, et al., Global waves synchronize the brain’s functional systems with fluctuating arousal. Sci Adv 7, eabf2709 (2021).

11. T. P. Lillicrap, A. Santoro, L. Marris, C. J. Akerman, G. Hinton, Backpropagation and the brain. Nat Rev Neurosci 21, 335–346 (2020).

12. D. E. Rumelhart, G. E. Hinton, R. J. Williams, Learning representations by back-propagating errors. Nature 323, 533–536 (1986).

13. X. Liu, D. A. Leopold, Y. Yang, Single-neuron firing cascades underlie global spontaneous brain events. Proceedings of the National Academy of Sciences 118, e2105395118 (2021).

14. Y. Yang, D. A. Leopold, J. H. Duyn, X. Liu, Hippocampal replay sequence governed by spontaneous brain-wide dynamics. PNAS Nexus pgae078 (2024). 10.1093/pnasnexus/pgae078.

15. Y. Yang, D. A. Leopold, J. H. Duyn, G. O. Sipe, X. Liu, Intrinsic forebrain arousal dynamics governs sensory stimulus encoding. bioRxiv (2023). 10.1101/2023.10.04.560900.

16. Y. Gu, et al., An orderly sequence of autonomic and neural events at transient arousal changes. Neuroimage 264, 119720 (2022).

17. M. D. Fox, et al., The human brain is intrinsically organized into dynamic, anticorrelated functional networks. Proceedings of the National Academy of Sciences 102, 9673–9678 (2005).

18. A. Mitra, A. Z. Snyder, T. Blazey, M. E. Raichle, Lag threads organize the brain’s intrinsic activity. Proceedings of the National Academy of Sciences 112, E2235–E2244 (2015).

19. T. Bolt, et al., A parsimonious description of global functional brain organization in three spatiotemporal patterns. Nat Neurosci 25, 1093–1103 (2022).

20. N. K. Logothetis, et al., Hippocampal–cortical interaction during periods of subcortical silence. Nature 491, 547–553 (2012).

21. G. Buzsáki, Hippocampal sharp wave-ripple: A cognitive biomarker for episodic memory and planning. Hippocampus 25, 1073–1188 (2015).

22. D. S. Margulies, et al., Situating the default-mode network along a principal gradient of macroscale cortical organization. Proceedings of the National Academy of Sciences 113, 12574–12579 (2016).

23. R. E. Yoss, N. J. Moyer, R. W. Hollenhorst, Pupil size and spontaneous pupillary waves associated with alertness, drowsiness, and sleep. Neurology 20, 545 (1970).

24. J. Reimer, et al., Pupil Fluctuations Track Fast Switching of Cortical States during Quiet Wakefulness. Neuron 84, 355–362 (2014).

25. J. H. Siegle, et al., Survey of spiking in the mouse visual system reveals functional hierarchy. Nature 592, 86–92 (2021).

26. S. E. J. de Vries, et al., A large-scale standardized physiological survey reveals functional organization of the mouse visual cortex. Nat Neurosci 23, 138–151 (2020).

27. N. A. Steinmetz, P. Zatka-Haas, M. Carandini, K. D. Harris, Distributed coding of choice, action and engagement across the mouse brain. Nature 576, 266–273 (2019).

28. D. C. Van Essen, et al., The Human Connectome Project: A data acquisition perspective. Neuroimage 62, 2222–2231 (2012).

29. W. Li, L. Zhu, L. Wen, Y. Yang, DeCap: Decoding CLIP Latents for Zero-Shot Captioning via Text-Only Training in The Eleventh International Conference on Learning Representations, (2023).

30. E. J. Allen, et al., A massive 7T fMRI dataset to bridge cognitive neuroscience and artificial intelligence. Nat Neurosci 25, 116–126 (2022).

31. A. Radford, et al., Learning Transferable Visual Models From Natural Language Supervision in Proceedings of the 38th International Conference on Machine Learning, Proceedings of Machine Learning Research., M. Meila, T. Zhang, Eds. (PMLR, 2021), pp. 8748–8763.

32. T. Chen, S. Kornblith, M. Norouzi, G. Hinton, A Simple Framework for Contrastive Learning of Visual Representations in Proceedings of the 37th International Conference on Machine Learning, Proceedings of Machine Learning Research., H. D. III, A. Singh, Eds. (PMLR, 2020), pp. 1597–1607.

33. L. van der Maaten, G. Hinton, Visualizing Data using t-SNE. Journal of Machine Learning Research 9, 2579–2605 (2008).

34. C. Higgins, et al., Replay bursts in humans coincide with activation of the default mode and parietal alpha networks. Neuron 109, 882–893.e7 (2021).

35. Y. Norman, O. Raccah, S. Liu, J. Parvizi, R. Malach, Hippocampal ripples and their coordinated dialogue with the default mode network during recent and remote recollection. Neuron 109, 2767–2780.e5 (2021).

36. A. Schaefer, et al., Local-Global Parcellation of the Human Cerebral Cortex from Intrinsic Functional Connectivity MRI. Cerebral Cortex 28, 3095–3114 (2018).

37. B. T. Thomas Yeo, et al., The organization of the human cerebral cortex estimated by intrinsic functional connectivity. J Neurophysiol 106, 1125–1165 (2011).

38. A. Arieli, A. Sterkin, A. Grinvald, A. Aertsen, Dynamics of Ongoing Activity: Explanation of the Large Variability in Evoked Cortical Responses. Science (1979) 273, 1868–1871 (1996).

39. M. S. Livingstone, D. H. Hubel, Effects of sleep and arousal on the processing of visual information in the cat. Nature 291, 554–561 (1981).

40. A. Hasenstaub, R. N. S. Sachdev, D. A. McCormick, State Changes Rapidly Modulate Cortical Neuronal Responsiveness. Journal of Neuroscience 27, 9607–9622 (2007).

41. C. Stringer, et al., Spontaneous behaviors drive multidimensional, brainwide activity. Science (1979) 364, eaav7893 (2019).

42. M. D. Fox, A. Z. Snyder, J. M. Zacks, M. E. Raichle, Coherent spontaneous activity accounts for trial-to-trial variability in human evoked brain responses. Nat Neurosci 9, 23–25 (2006).

43. B. J. He, Spontaneous and Task-Evoked Brain Activity Negatively Interact. Journal of Neuroscience 33, 4672–4682 (2013).

44. M. J. McGinley, et al., Waking State: Rapid Variations Modulate Neural and Behavioral Responses. Neuron 87, 1143–1161 (2015).

45. W. Chen, K. Park, Y. Pan, A. P. Koretsky, C. Du, Interactions between stimuli-evoked cortical activity and spontaneous low frequency oscillations measured with neuronal calcium. Neuroimage 210, 116554 (2020).

46. M. A. Wilson, B. L. McNaughton, Reactivation of Hippocampal Ensemble Memories During Sleep. Science (1979) 265, 676–679 (1994).

47. D. R. Euston, M. Tatsuno, B. L. McNaughton, Fast-Forward Playback of Recent Memory Sequences in Prefrontal Cortex During Sleep. Science (1979) 318, 1147–1150 (2007).

48. Y. Norman, et al., Hippocampal sharp-wave ripples linked to visual episodic recollection in humans. Science (1979) 365, eaax1030 (2019).

49. A. P. Vaz, S. K. Inati, N. Brunel, K. A. Zaghloul, Coupled ripple oscillations between the medial temporal lobe and neocortex retrieve human memory. Science (1979) 363, 975–978 (2019).

50. J. J. Sakon, M. J. Kahana, Hippocampal ripples signal contextually mediated episodic recall. Proceedings of the National Academy of Sciences 119, e2201657119 (2022).

51. J. Karimi Abadchi, et al., Spatiotemporal patterns of neocortical activity around hippocampal sharp-wave ripples. Elife 9, e51972 (2020).

52. R. Kaplan, et al., Hippocampal Sharp-Wave Ripples Influence Selective Activation of the Default Mode Network. Current Biology 26, 686–691 (2016).

53. J. G. Klinzing, N. Niethard, J. Born, Mechanisms of systems memory consolidation during sleep. Nat Neurosci 22, 1598–1610 (2019).

54. Y. Dudai, A. Karni, J. Born, The Consolidation and Transformation of Memory. Neuron 88, 20–32 (2015).

55. J. Bayer, J. Gläscher, J. Finsterbusch, L. H. Schulte, T. Sommer, Linear and inverted U-shaped dose-response functions describe estrogen effects on hippocampal activity in young women. Nat Commun 9, 1220 (2018).

56. E. Baldi, C. Bucherelli, The Inverted “U-Shaped” Dose-Effect Relationships in Learning and Memory: Modulation of Arousal and Consolidation. Nonlinearity Biol Toxicol Med 3, nonlin.003.01.002 (2005).

57. G. Zerbes, F. M. Kausche, J. C. Müller, K. Wiedemann, L. Schwabe, Glucocorticoids, Noradrenergic Arousal, and the Control of Memory Retrieval. J Cogn Neurosci 31, 288–298 (2019).

58. D. J.-F. de Quervain, B. Roozendaal, J. L. McGaugh, Stress and glucocorticoids impair retrieval of long-term spatial memory. Nature 394, 787–790 (1998).

59. X. Liu, et al., Modulation of cross-hierarchy propagating waves across sleep stages in ISMRM & ISMRT Annual Meeting, (2024), p. 8360.

60. A. G. Siapas, M. A. Wilson, Coordinated Interactions between Hippocampal Ripples and Cortical Spindles during Slow-Wave Sleep. Neuron 21, 1123–1128 (1998).

61. S. Diekelmann, J. Born, The memory function of sleep. Nat Rev Neurosci 11, 114–126 (2010).

62. S. Lecci, et al., Coordinated infraslow neural and cardiac oscillations mark fragility and offline periods in mammalian sleep. Sci Adv 3, e1602026 (2017).

63. C. Kjaerby, et al., Memory-enhancing properties of sleep depend on the oscillatory amplitude of norepinephrine. Nat Neurosci 25, 1059–1070 (2022).

64. J. M. Shine, et al., The Dynamics of Functional Brain Networks: Integrated Network States during Cognitive Task Performance. Neuron 92, 544–554 (2016).

65. D. S. Bassett, et al., Dynamic reconfiguration of human brain networks during learning. Proceedings of the National Academy of Sciences 108, 7641–7646 (2011).

66. U. Braun, et al., Dynamic reconfiguration of frontal brain networks during executive cognition in humans. Proceedings of the National Academy of Sciences 112, 11678–11683 (2015).

67. W. Huijbers, et al., Explaining the encoding/retrieval flip: Memory-related deactivations and activations in the posteromedial cortex. Neuropsychologia 50, 3764–3774 (2012).

68. S. Daselaar, et al., Posterior midline and ventral parietal activity is associated with retrieval success and encoding failure. Front Hum Neurosci 3 (2009).

69. P. Vannini, et al., What Goes Down Must Come Up: Role of the Posteromedial Cortices in Encoding and Retrieval. Cerebral Cortex 21, 22–34 (2011).

70. F. Han, et al., Reduced coupling between cerebrospinal fluid flow and global brain activity is linked to Alzheimer disease–related pathology. PLoS Biol 19, e3001233.(2021).

71. F. Han, X. Liu, R. B. Mailman, X. Huang, X. Liu, Resting-state global brain activity affects early β-amyloid accumulation in default mode network. Nat Commun 14, 7788 (2023).

72. F. Han, et al., Reduced coupling between cerebrospinal fluid flow and global brain activity is linked to tau pathology. Alzheimer’s & Dementia 19, e075860 (2023).

73. J. J. Iliff, et al., Cerebral Arterial Pulsation Drives Paravascular CSFInterstitial Fluid Exchange in the Murine Brain. Journal of Neuroscience 33, 18190–18199 (2013).

74. S. J. van Veluw, et al., Vasomotion as a Driving Force for Paravascular Clearance in the Awake Mouse Brain. Neuron 105, 549–561.e5 (2020).

75. L. Xie, et al., Sleep Drives Metabolite Clearance from the Adult Brain. Science (1979) 342, 373–377 (2013).

76. P. J. Whitehouse, et al., Alzheimer’s Disease and Senile Dementia: Loss of Neurons in the Basal Forebrain. Science (1979) 215, 1237–1239 (1982).

77. R. T. Bartus, On Neurodegenerative Diseases, Models, and Treatment Strategies: Lessons Learned and Lessons Forgotten a Generation Following the Cholinergic Hypothesis. Exp Neurol 163, 495–529 (2000).

78. D. A. Drachman, J. Leavitt, Human Memory and the Cholinergic System: A Relationship to Aging? Arch Neurol 30, 113–121 (1974).

79. M. E. Hasselmo, The role of acetylcholine in learning and memory. Curr Opin Neurobiol 16, 710–715 (2006).

