## Supplementary Materials for "Propagating cortical waves coordinate sensory encoding and memory retrieval in the human brain"

3.2.2 Allen Institute Visual Coding Mice Two-photon Calcium Imaging Dataset 15

### 1 Supplementary Tables

#### 1.1 Table S1

**Table S1**. List of thalamic nuclei and their abbreviation as in the Morel atlas [1].

| Abbreviation | Full Name |
|---|---|
| AD | anterior dorsal nucleus |
| AM | anterior medial nucleus |
| AV | anterior ventral nucleus |
| LD | lateral dorsal nucleus |
| VAmc | ventral anterior nucleus magnocellular part |
| VApc | ventral anterior nucleus parvocellular part |
| VLa | ventral lateral anterior nucleus |
| VLpd | ventral lateral posterior nucleus dorsal part |
| VLpv | ventral lateral posterior nucleus ventral part |
| VM | ventral medial nucleus |
| VPI | ventral posterior inferior nucleus |
| VPLa | ventral posterior lateral nucleus anterior part |
| VPLp | ventral posterior lateral nucleus posterior part |
| VPM | ventral posterior medial nucleus |
| LGNmc | Lateral geniculate nucleus magnocellular part |
| LGNpc | Lateral geniculate nucleus parvocellular part |
| MGN | Medial geniculate nucleus |
| Po | posterior nucleus |
| SG | suprageniculate nucleus |
| LP | lateral posterior nucleus |
| PuA | anterior pulvinar nucleus |
| PuI | inferior pulvinar nucleus |
| PuL | lateral pulvinar nucleus |
| PuM | medial pulvinar nucleus |
| Hb | habenular nucleus |
| CeM | central median nucleus |
| MV | medioventral nucleus |
| Pv | paraventricular nucleus |
| CL | central lateral nucleus |
| CM | centre median nucleus |
| Pf | parafascicular nucleus |
| sPf | subparafascicular nucleus |
| MDmc | mediodorsal nucleus magnocellular part |
| MDpc | mediodorsal nucleus parvocellular part |

#### 1.2 Table S2

**Table S2**. List of brain regions and their abbreviation as in the AAN atlas [2].

| Abbreviation | Full Name |
|---|---|
| DR | dorsal raphe |
| LC | locus coeruleus |
| MRF | midbrain reticular formation |
| MR | median raphe |
| PAG | periaqueductal gray |
| PBC | parabrachial complex |
| PO | pontis oralis |
| PPN | pedunculopontine nucleus |
| VTA | ventral tegmental area |

### 2 Supplementary Figures

#### 2.1 Figure S1

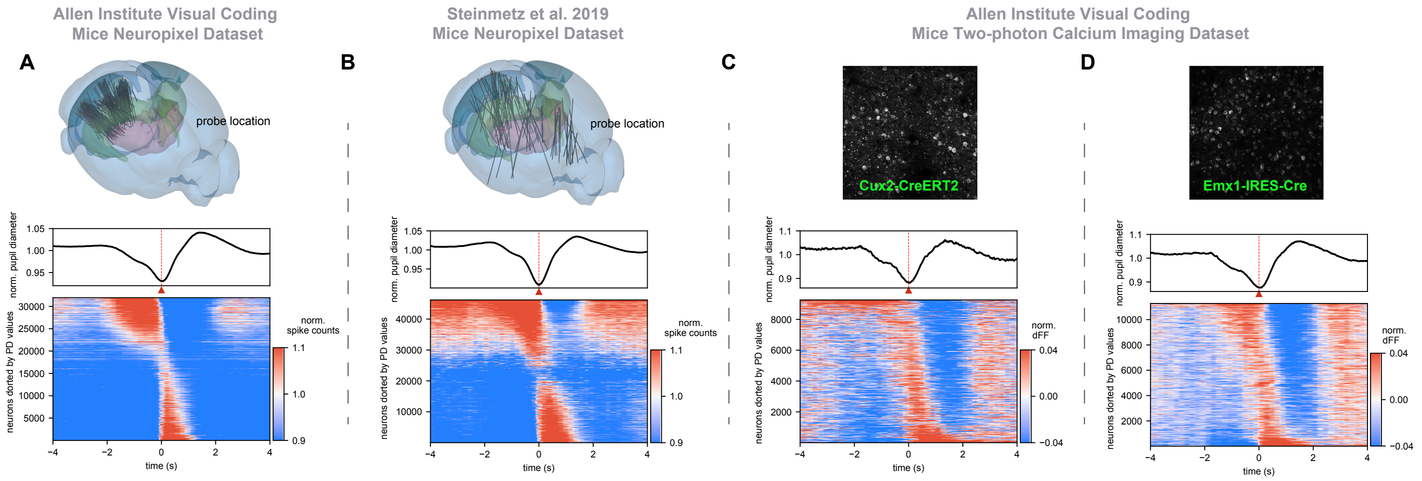

**Figure S1. Pupil dilation is accompanied by stereotypical infra-slow, brain-wide sequential neural activation, evident across different cell types and extending through various brain regions in mice.** (A) Analysis of Allen Institute Visual Coding Neuropixel dataset. Top row: visualization of 3-dimensional probes location in Allen Mouse Brain Common Coordinate Framework (version 3). Second row: The normalized pupil diameter (Top) and neuronal spiking activities (Bottom), both averaged around the onset of pupil dilation over an 8-second time window from all N mice, with neurons sorted by the principal delay profile identified during periods of pupil dilation. Similar analytical approaches are applied to another mice neuropixel dataset with probes recording covering the whole brain (B), and to two-photon calcium imaging datasets of two types of excitatory neurons in mice visual regions, i.e. Cux2-CreERT2 (C) and Emx1-IRES-Cre (D).

#### 2.2 Figure S2

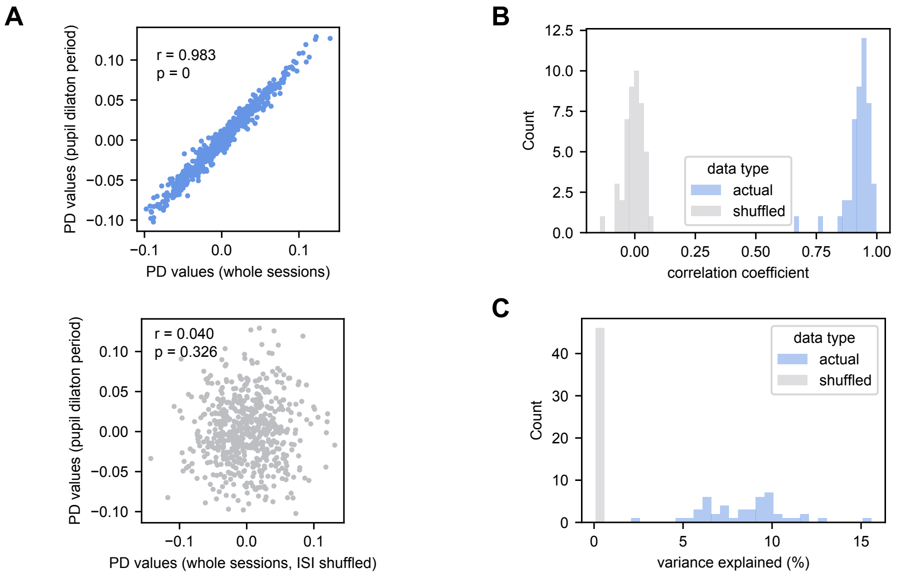

**Figure S2. The sequential activation during pupil dilation is the principal infra-slow dynamics in mice brain.** (A) High degree of similarity between the principal delay profile derived from periods of pupil dilation and that from whole recording sessions in an example mouse (Top). This correlation significantly diminishes when neural activities are randomized by inter-spike interval (ISI) permutation (Bottom). (B) The distribution of similarity, quantified by Pearson's correlation coefficient, between the principal delay profiles derived from pupil dilation periods and from whole recording sessions across all studied mice, showing high degree of similarity generalizable to all the mice. (C) Distribution of the variance explained by the principal delay profile derived from whole session data for neural activities during pupil dilation periods. This analysis validates the high degree of similarity between the infra-slow dynamics characterizing the entire recording session and those specifically associated with pupil dilation periods.

#### 2.3 Figure S3

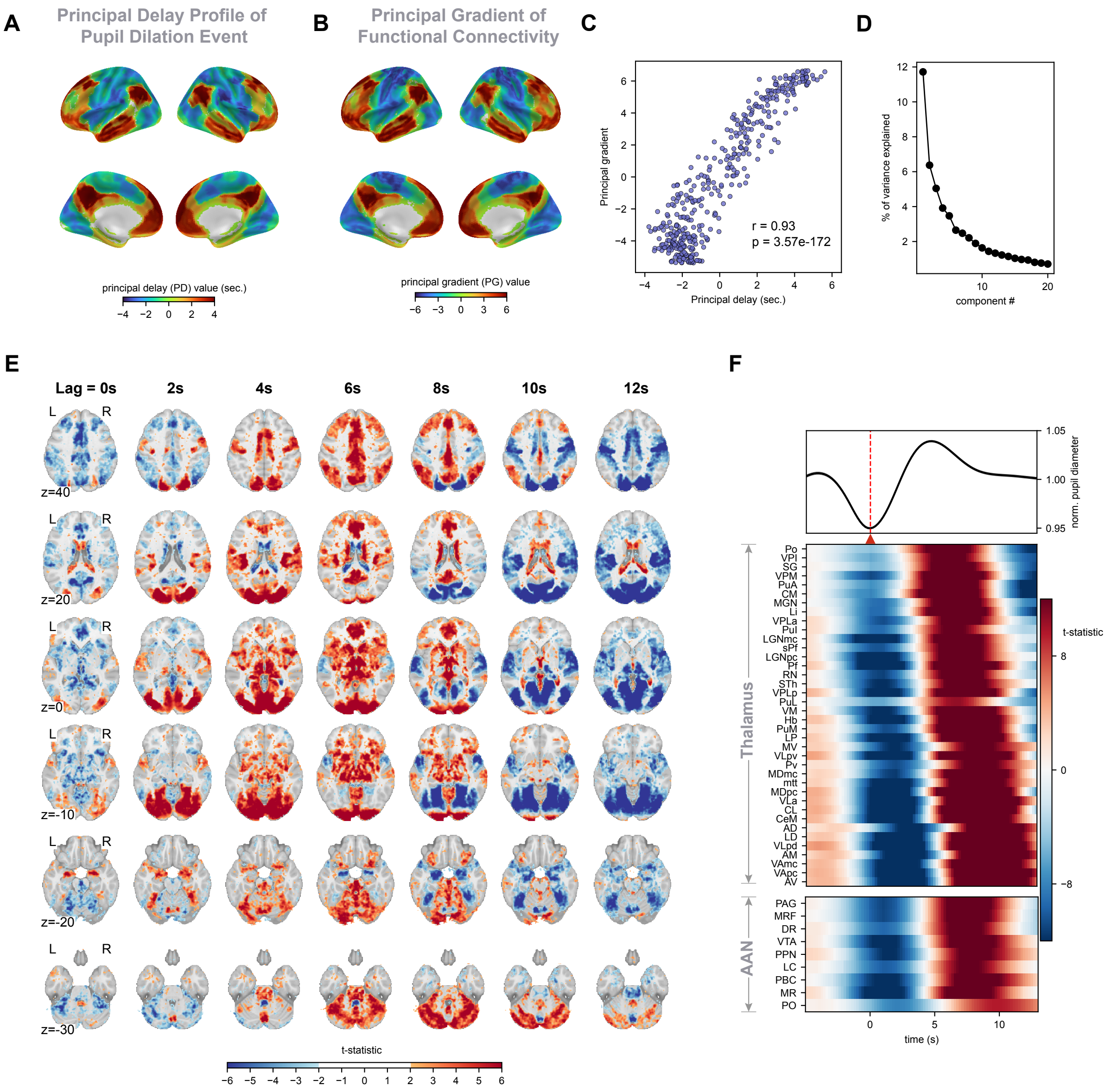

**Figure S3. Resting-state pupil dilation is accompanied by hierarchical propagating waves in human brain.** (A) The principal delay (PD) profile derived for the period of pupil dilation quantifying the temporal shift of each brain region in relation to the onset of pupil dilation. (B) The principal gradient (PG) of functional connectivity, characterizing the topographical organization of the cortex with regions at one end serving sensory/motor (SM) functions and the other end known as the default-mode network (DMN). (C) Strong similarity between the PD profile (shown in A) and PG (shown in B) and quantified using Pearson's correlation coefficient. (D) The scree plot showing the percentage of variance explained by the first 20 prinicpal components derived during periods of pupil dilation. (E) The BOLD signal statistical map shown for 7 evenly spaced time lags, from 0 to 12 seconds following the onset of pupil dilation, with each row showing a distinct brain slice. (F) The temporal dynamics of pupil size (Top) and subcortical regions within the temporal window surrounding pupil dilation onset. The thalamic regions of interest (ROIs) are defined according to the Morel's atlas (Middle) and 9 brainstem ROIs are defined by the Harvard AAS atlas (Bottom). The full name of the abbreviation can be found in Table S1 and Table S2.

#### 2.4 Figure S4

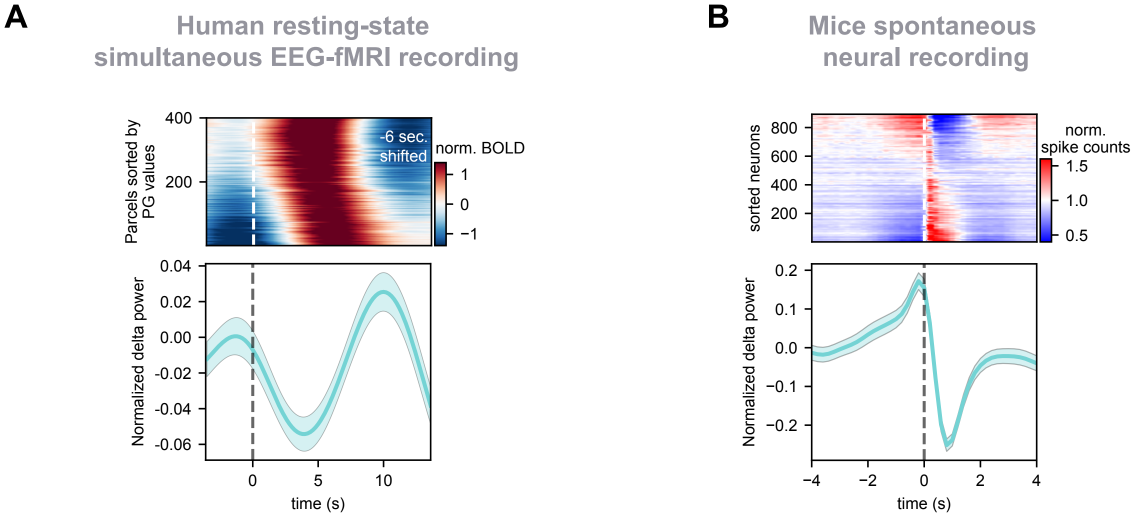

**Figure S4. Association between brain-wide sequential activity and arousal modulation in humans and mice.** (A) Cortical fMRI ROI signals (Top) and EEG delta-band power (Bottom) are averaged according to the detected SM-DMN propagation center across all resting-state simultaneous EEG-fMRI recording sessions. The averaged propagation pattern is adjusted by a 6-second left shift to account for hemodynamic delay, with a white dashed line marking time zero to indicate the onset of an increasing global signal. (D) Spiking activity (Top) and delta-band power (Bottom) are averaged according to the time of positve-delay neuron onset within the detected spiking cascade across all mice from Allen Institute Visual Coding Neuropixel dataset. The neuronal spiking data are organized in the form of spiking cascade by sorting the neurons according to the principal delay profile. Together, these findings highlights the unified coordination between stereotypical brain-wide neural activity and arousal modulation across species.

#### 2.5 Figure S5

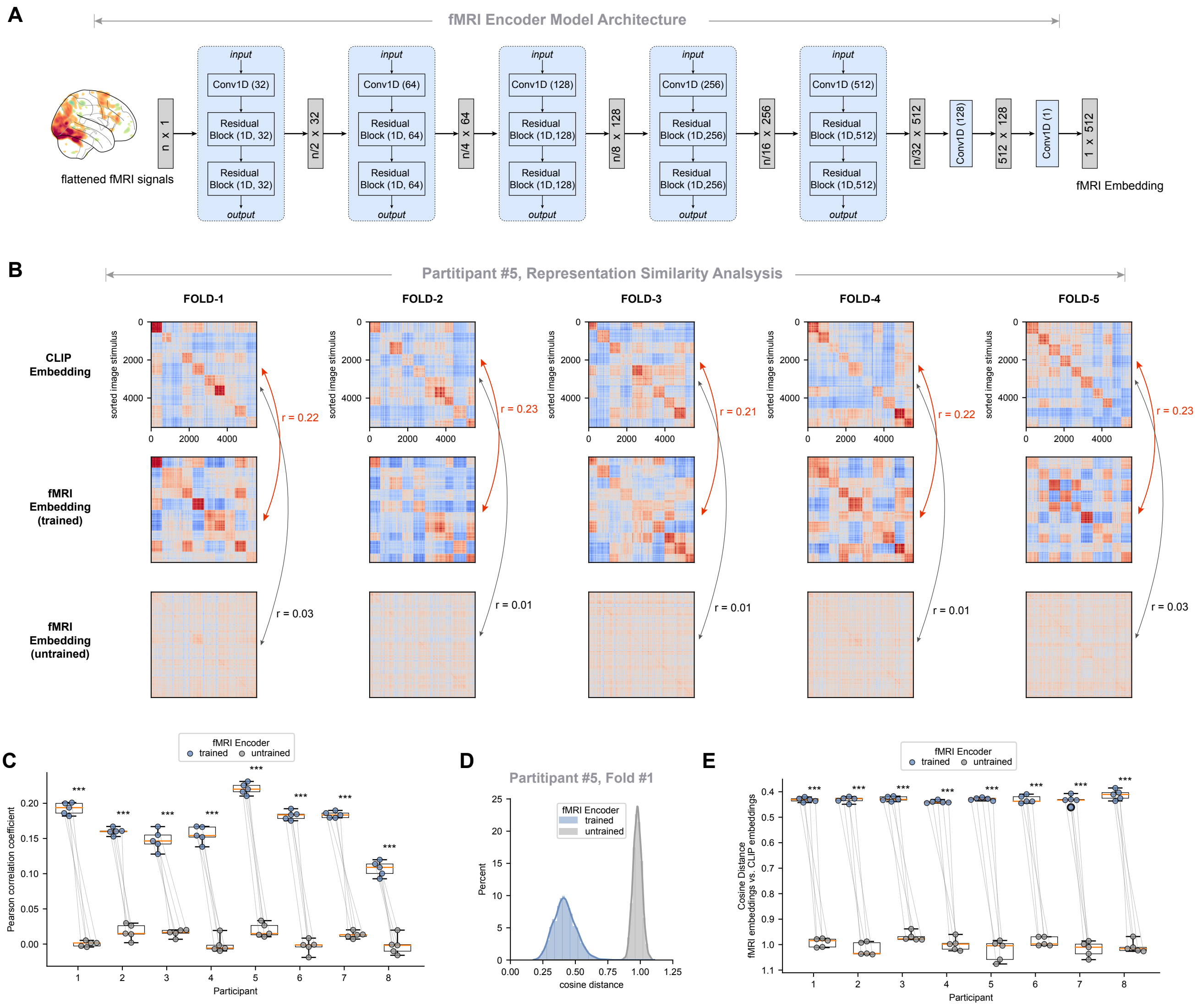

**Figure S5. Bridging fMRI and CLIP embedding spaces through a deep learning-based fMRI encoder.** (A) The model architecture of fMRI encoder, containing several convolutional layers to transform the fMRI BOLD response to 512-dimentional fMRI embedding vector, i.e., the latent representation of fMRI signal. (B) Representation similarity analysis for an example participant, with fMRI encoders are trained and evaluated with 5-fold cross-validation. For each validation fold (a column in the panel), representation dissimilarity matrices (RDMs) are constructed for semantic CLIP embeddings (Top), fMRI embeddings extracted with trained fMRI encoder (Middle), and fMRI embeddings extracted with untrained fMRI encoder (Bottom). For the RDMs within each fold, image stimuli are sorted identically according to CLIP-derived RDMs. Notably, RDMs based on fMRI embeddings show a significant improvement in similarity to CLIP-derived RDMs when extracted by the trained encoder (indicated by red arrows), compared to the untrained encoder (black arrows). (C) Boxplots comparing the correlation differences between CLIP RDMs and RDMs from trained versus untrained fMRI embeddings across subjects. Each dot represents a validation fold from an individual subject, with pair-wise t-test (two-sided, N=5) used for statistical significance test. (D) Distribution of cosine distance, measuring the discrepancy between CLIP embeddings and fMRI embeddings, summarized separately for trained and untrained fMRI encoders from an example validation fold. (E) Boxplots showing the significant decrease in cosine distance (i.e., discrepancy) between CLIP and fMRI embeddings following the training of the fMRI encoder.

#### 2.6 Figure S6

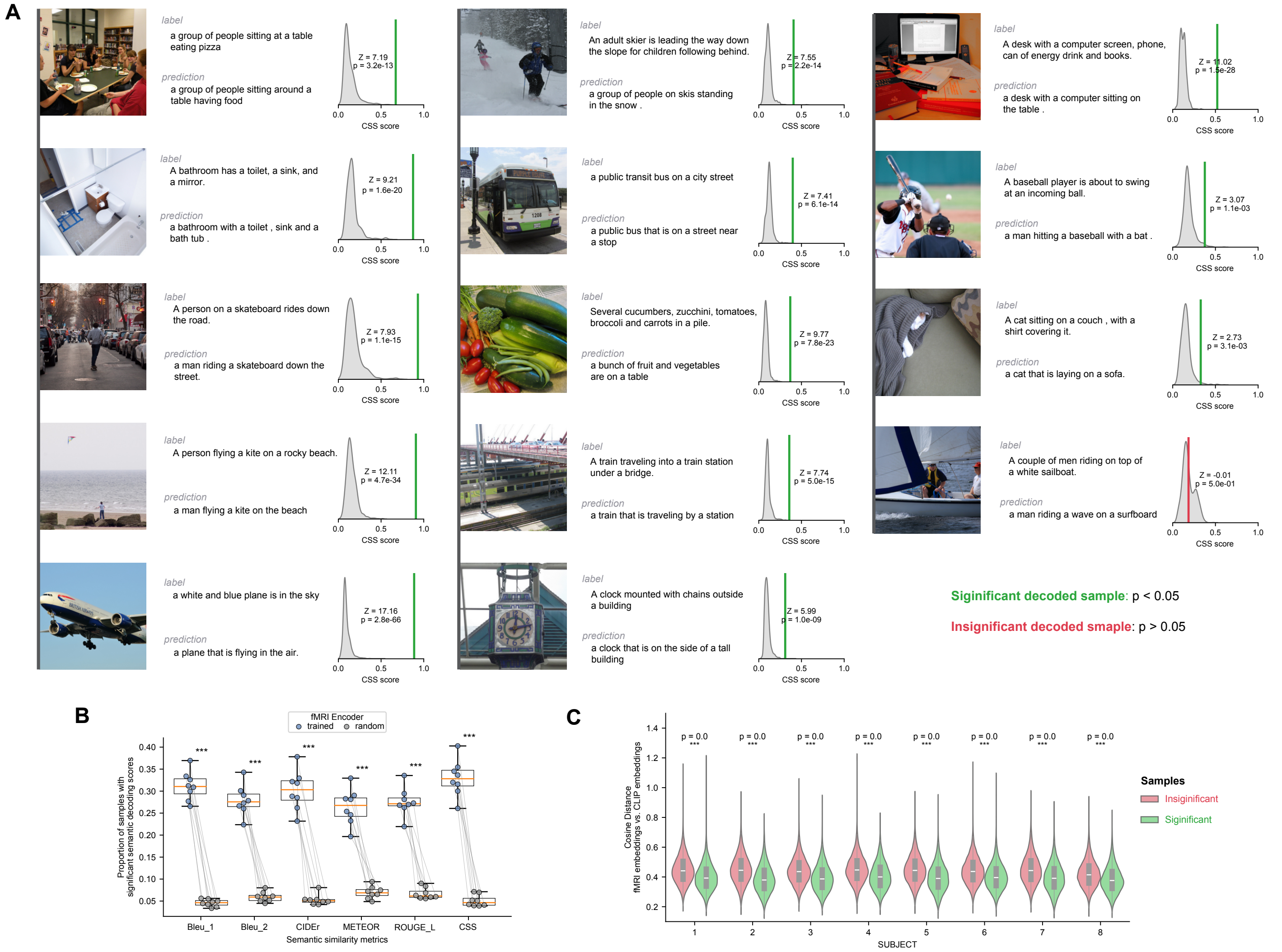

**Figure S6. Decoding semantic information from stimuli-evoked BOLD response.** (A) Examples of significantly and insignificantly decoded trials. Each trail includes the image stimulus presented to human participant (Left), alongside the model-generated caption and the human-labeled ground truth caption (Middle), with their similarity quantified using composite semantic similarity (CSS) scores. To assess the statistical significance of the decoding, for each trial, the actual CSS score is compared against a null distribution, constructed based on CSS scores between the ground truth caption and 1,000 randomly generated captions by sampling the CLIP embedding space. Notably, the null distribution varies across trials, reflecting differences in decoding complexity tied to the semantic content of the stimuli. Trials are considered significantly decoded if their p-value is below 0.05, indicated by green. (B) Box plot comparing encoding accuracy—quantified as the proportion of significantly decoded trials—between the trained and untrained (control) fMRI encoder. Results are presented for various semantic similarity metrics, with each dot representing an individual participant. Statistical significance is evaluated using a pairwise t-test (two-sided, N=8). (C) Violin plot showing the distribution of the cosine distance between the fMRI embeddings and CLIP embeddings, comparing between the insignificant (red) and significant (green) decoded trials, summarized for each of the participants. Aligning with expectations, trials that are significantly decoded exhibit a closer alignment (reduced cosine distance) between the fMRI embeddings and the CLIP embeddings.

#### 2.7 Figure S7

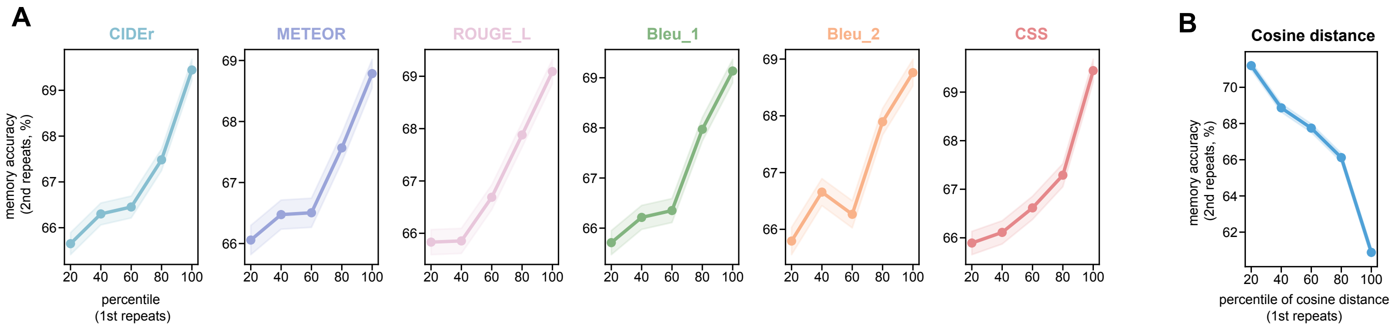

**Figure S7. Semantic encoding accuracy of first-time image presentation predicts subsequent memory recall.** (A) Influence of first-time encoding accuracy on subsequent memory task performance. Stimuli are categorized based on the percentile of their initial encoding accuracy, with increments of 20%, and the average memory recall accuracy during their second presentation is calculated for each bin. Each column shows the results for specific metrics to evaluate the semantic similarity score, from left to right showing CIDEr, METEOR, ROUGE_L, Bleu_1, Bleu_2, and CSS scores. (B) Parallel analysis to (A) but with a focus on the effect of the initial alignment between fMRI and CLIP embeddings (measured by cosine distance) on subsequent memory performance. A closer initial alignment (reduced cosine distance), indicative of better sensory encoding within the brain signal, is shown to enhance memory recall.

#### 2.8 Figure S8

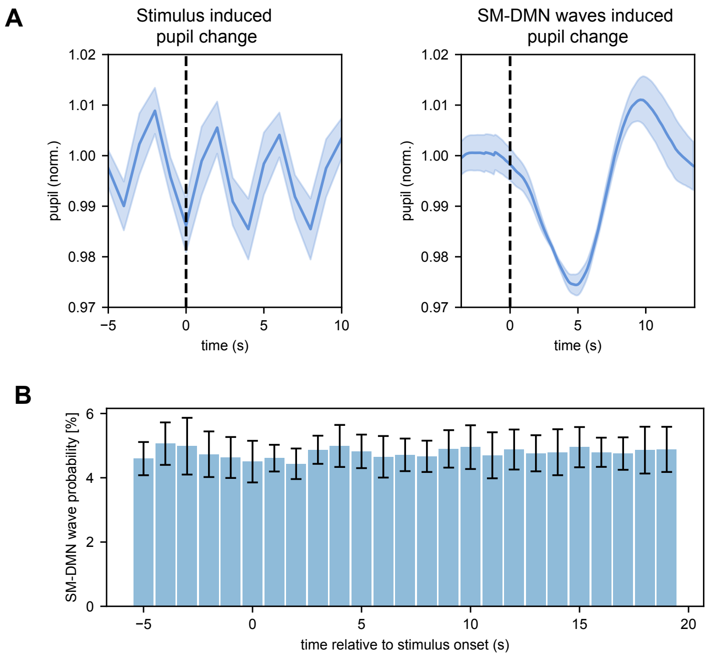

**Figure S8. Influence of external stimulation to pupil dynamics and SM-to-DMN wave occurrences**. (A) Left: The averaged pupil fluctuation across the onsets of stimuli with time zero defined as the timing of stimulus onset. Right: The averaged pupil size across detected SM-to-DMN waves with time zero defined as the onset of global mean signal increase. The shaded area indicates the standard error of the mean (SEM). (C) The histogram showing the probability of occurrence for SM-to-DMN waves, with time zero set at the onset of the stimulus. To avoid overlapping between segments for averaging, stimuli were selected with a minimum interval of 40 seconds between each. We also excluded the stimuli that within first 40 seconds and last 40 seconds of fMRI scans to mitigate edge effects. Error bars represent the standard deviation.

#### 2.9 Figure S9

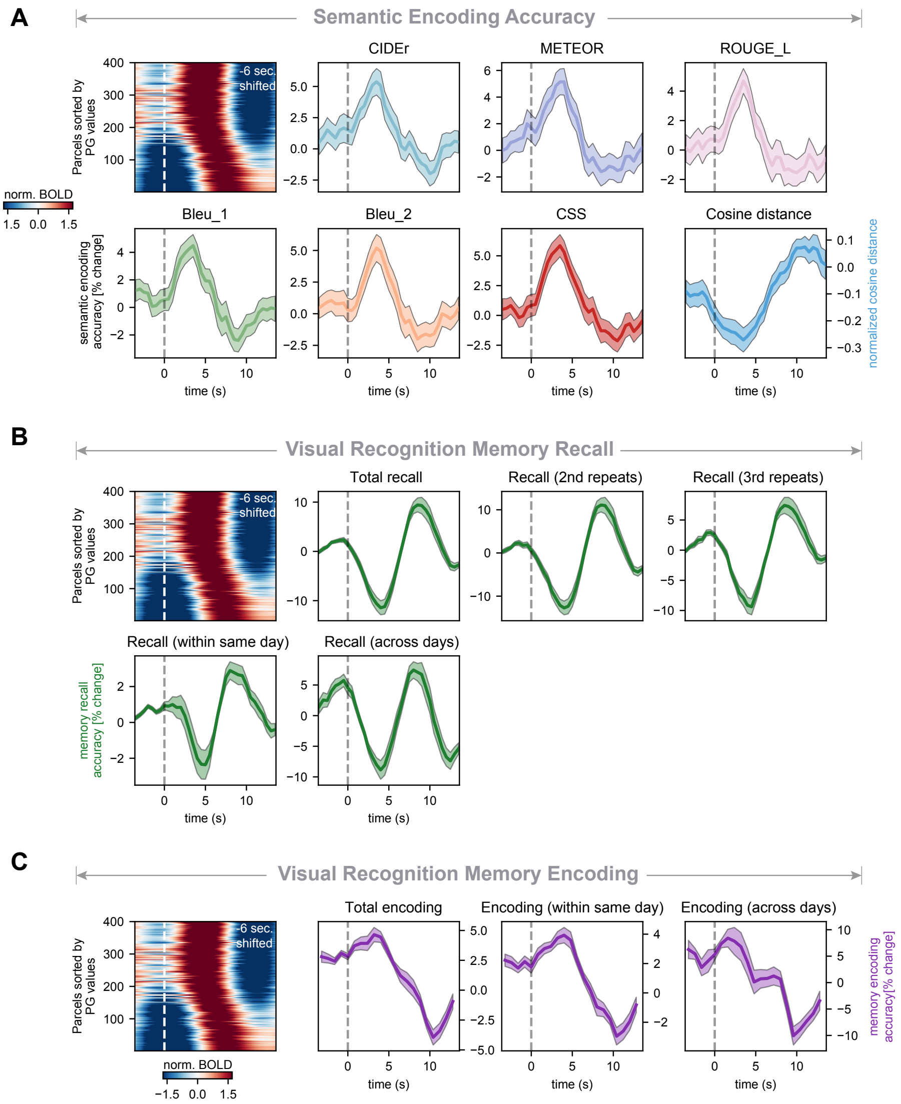

**Figure S9. Sensory encoding and memory recall are oppositely modulated across the cycle of the SM-to-DMN propagating wave.** (A) Phase-dependent modulation of sensory processing efficacy, quantified through semantic encoding accuracy using various semantic similarity metrics (CIDEr, METEOR, ROUGE_L, Bleu_1, Bleu_2, and CSS). The cosine distance measures the alignment between fMRI and CLIP embeddings, with closer alginment (reduced cosine distance) indicating better sensory encoding quality within the brain signal. Consistently, the sensory processing shows enhanced encoding efficacy during the early phase of propagation cycle. (B) Phase-dependent modulation of memory recall. Various recall metrics are incorporated, including total recall (the combined recall from second and third image presentations), recalls specific to the second and third presentations, short-term recall (recalls derived for the scenario where first and second image presented within the same day), and recall across different days. Consistently, memory recalls show significant enhancement during the latter phase of the propagation cycle. (C) Phase-dependent modulation of memory encoding. Various encoding metrics are considered, including total encoding accuracy (across all first and second presentation pairs), short-term encoding (all encoding within the same day), and encoding across different days. Consistently, memory encoding are significantly improved during the early phase of the propagation cycle. The averaged pattern of the detected SM-DMN propagation left shifted by 6 seconds, with time zero marked by white dashed line indicating the onset of increasing global signal.

#### 2.10 Figure S10

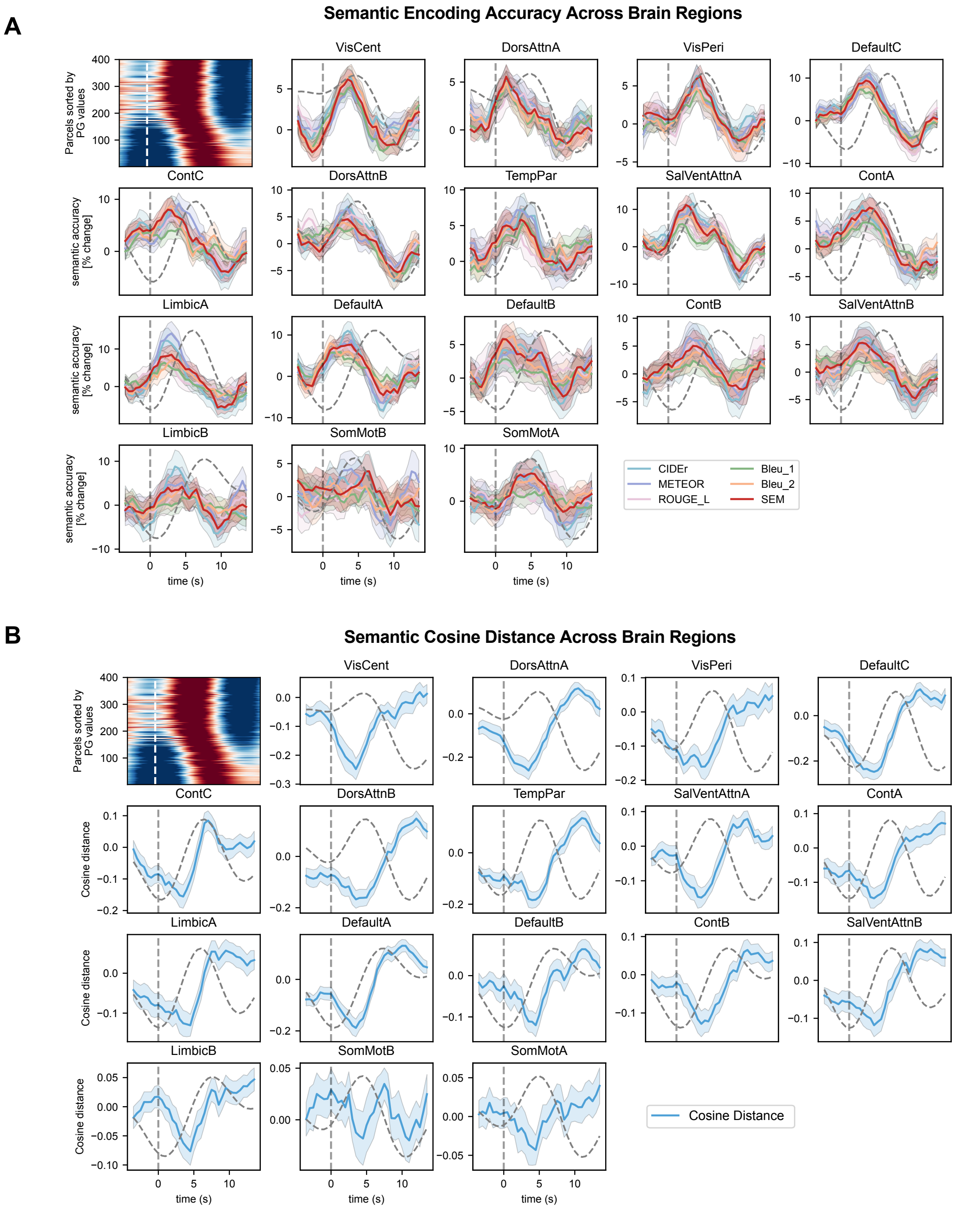

**Figure S10. Regional dynamics of sensory encoding efficacy during SM-to-DMN propagation.** (A) Semantic encoding accuracy evaluated with various semantic similarity metrics (CIDEr, METEOR, ROUGE_L, Bleu_1, Bleu_2, and CSS). (B) Modulation of cosine distance, which measures the alignment between fMRI and CLIP embeddings (reduced cosine distance indicating better sensory encoding quality), consistently aligns with the modulation of encoding accuracy in (A). For both panels, the regions are organized in descending order according to the significance of their semantic encoding accuracy, revealing a gradient in the modulation effect. The averaged propagation pattern is adjusted by a 6-second left shift to account for the hemodynamic delay, with a white dashed line marking time zero to indicate the onset of an increasing global signal. For comparison, the average activation for each region is marked with gray dotted line, which is shifted ahead of time by 6 seconds to account for the hemodynamic response delay.

### 3 Materials and Methods

#### 3.1 Human fMRI Datasets

##### 3.1.1 Human Connectome Project

We utilized the WU-UMinn Human Connectome Project (HCP) 7T dataset, a subset of the HCP S1200 release [3], comprising 7T fMRI data from 184 subjects within the age range of 22 to 35. Our analysis focused on two 15-minute, eyes-open resting-state fMRI sessions, with repetition time (TR) of 1s and 1.6 mm isotropic voxels. Simultaneous eye tracking was conducted using an EyeLink device with a sampling rate of 1000 Hz. Further details about the HCP dataset can be found in [3].

##### 3.1.2 Natural Scenes Dataset

The Natural Scenes Dataset (NSD) includes whole-brain 7T fMRI scans with a repetition time (TR) of 1.6s and 1.8mm isotropic voxels, conducted during a visual memory task. This dataset involved eight meticulously selected human participants who were shown between 9000 and 10000 natural scene images. Each image was presented three times, resulting in a total of 22000 to 30000 trials across a span of one year. For every trial, as participants viewed an image stimulus, they were required to report whether they perceived the image as novel. Additional details about the dataset can be found in [4].

##### 3.1.3 Simultaneous EEG-fMRI Resting-state Dataset

Resting-state fMRI data were collected simultaneously with electroencephalography (EEG) for 27 subjects (average age: $22.1 \pm 3.1$ years, including 14 females). For our analysis, we only utilized the 10-minute resting-state scan. The fMRI imaging data were acquired using a 3T scanner, with a repetition time (TR) of 2.1s and 3mm isotropic voxels. The EEG data were gathered using a 32-channel MR-compatible EEG system, with a recording sampling rate of 5000Hz. Additional details can be found in [5].

#### 3.2 Mice Single-Neuron Recording Datasets

##### 3.2.1 Allen Institute Visual Coding Mice Neuropixel Dataset

The Allen Institute Visual Coding Neuropixel dataset comprises high-density extracellular neuron recordings of mice using Neuropixel probes [6, 7]. Each mouse was implanted up to six Neuropixel probes, which targeted the primary visual cortex (VISp) and five high-order visual cortical areas, namely, latero-medial area (VISl), anterol-ateral area (VISal), rostro-lateral area (VISrl), postero-medial area (VISpm), and antero-medial area (VISam). The silicon probes were inserted to a depth of up to 3.5mm into the brain, enabling the recording of spiking activity within two visual thalamic nuclei, i.e., the lateral posterior nucleus (LP) and the lateral geniculate nucleus (LGN), as well as other regions that the probes traversed, such as the hippocampus.

##### 3.2.2 Allen Institute Visual Coding Mice Two-photon Calcium Imaging Dataset

The Allen Institute Visual Coding Mice Two-photon Calcium Imaging Dataset comprises single-neuron recordings from the mouse visual cortex, obtained through 2-photon fluorescence imaging. Utilizing transgenic tools, these recordings specifically targeted the activities of distinct populations of Cre-defined neurons. The dataset includes a total of 63,251 neurons from 14 different transgenic lines, covering 6 cortical areas and 4 cortical layers. For our study, we focused on two Cre lines (Cux2-CreERT2 and Emx1-IRES-Cre) that had the highest number of neurons recorded. In addition to neural recordings, both running speed and pupil size were simultaneously monitored. Further details can be found in [7].

##### 3.2.3 UCL Brain-wide Mice Neuropixel Recording Dataset

The UCL mice dataset includes recordings from approximately 30,000 neurons across 43 brain regions in mice, utilizing Neuropixels probes to cover the entire left hemisphere. In each mouse, two or three probes were inserted simultaneously, allowing for the concurrent recording of hundreds of neurons during each session. The study comprised 92 probe insertions across 39 sessions from 10 mice, with an average of approximately 747 neurons recorded per session. Alongside neural activity, both running speed and pupil size were simultaneously monitored and recorded. Additional details are available in [8].

#### 3.3 fMRI Preprocessing

##### 3.3.1 HCP

The resting-state HCP fMRI volumetric data were preprocessed using the minimal preprocessing pipeline [9] and artifacts were further removed with ICA-FIX denoising [10]. fMRI data were then spatially smoothed with 2mm Full Width at Half Maximum (FWHM) Gaussian kernel and temporal filtering within a bandpass range of 0.001-0.15Hz. Finally, signal from each voxel was standardized by subtracting the mean and dividing by the standard deviation.

##### 3.3.2 NSD

The NSD fMRI data underwent initial preprocessing to correct for head motion, EPI distortion, gradient nonlinearities, and alignment across scan sessions [4]. Analyses of fMRI data for each subject were performed in the subject-native space.

For the fMRI time series analysis, i.e. propagation, nuisance parameters such as motion, white matter, and cerebrospinal fluid (CSF) signals were regressed out from the volumetric fMRI data, which were then smoothed spatially with 2mm FWHM Gaussian kernel and temporally with bandpass filtering of 0.001-0.15Hz. Each voxel's signal was then standardized to have zero mean and unit standard deviation.

For analyzing the hemodynamic response of single trials (inputs of the decoding models), GLMdenoise—a generalized linear model (GLM) approach was used to provide estimates of the BOLD amplitude while reducing noise by integrating nuisance regressor [4, 11].

##### 3.3.3 Simultaneous EEG-fMRI

The resting-state fMRI BOLD data was preprocessed using script from the 1000 Functional Connectomes Project with slight modification [12]. Nuisance parameters, including linear and quadratic trends, motion parameters, white matter, and CSF signals, were regressed from the fMRI data. The volumetric data then was smoothed spatially with 2mm FWHM Gaussian kernel, and temporally filtered with a bandpass range of 0.001-0.15Hz. The voxel's signal was finally standardized to have zero mean and unit standard deviation [5].

##### 3.3.4 Network Parcellations

For every fMRI dataset analyzed, we utilized the Schaefer 400 Parcellations (Yeo-17 Network version) [13] to obtain cortical signals from the volumetric fMRI data. This was accomplished by averaging the standardized voxel signals located within each parcel. Similarly, signals from thalamic nuclei/regions were extracted utilizing the Morel Atlas [1], and signals from the brainstem nuclei, part of the ascending arousal network, using the Harvard AAN atlas [2].

#### 3.4 Pupil Size

Pupil areas recorded for humans were obtained using the EyeLink device, with raw data sampled at 2000Hz for the NSD dataset and 1000Hz for the HCP dataset. Prior to downsampling the time series to 50Hz, we converted the pupil area into pupil diameters. Missing pupil data, resulting from false detections or eye blinks, were interpolated using data from the nearest time points. Subsequently, we synchronized the pupil data with each TR of the concurrent fMRI signal. Periods, where the pupil size data had significant missing information (more than 50%), were removed from subsequent analyses.

Pupil areas recorded for mice were captured using cameras, with a sampling rate of 100Hz for the UCL Neuropixel dataset and 30Hz for both the Allen Institute Neuropixel and two-photon datasets. Initially, we converted the pupil area into diameters and then resampled the data to a uniform rate of 30Hz. Following this, missing pupil data were interpolated using the nearest time points to ensure continuity in the dataset.

To identify dilation events within infra-slow pupil fluctuations, we utilized a low-pass filter on the pupil size time series, setting a cut-off frequency at 0.15Hz for human datasets and 0.3Hz for mouse datasets. Pupil dilation events were defined as the periods where dilation lasted for at least 1 second in the filtered pupil size data.

#### 3.5 EEG Preprocessing

The EEG data were preprocessed to remove the gradient and ballistocardiogram artifacts from each channel, utilizing algorithms detailed in [14]. Following this, the data were subjected to low-pass filtering with a cut-off frequency of 125Hz. Pulse artifacts were removed through independent component analysis (ICA), and signals were corrected for distortions to account for distortions attributable to head motion [15]. More comprehensive description of the EEG signal preprocessing can be found in [5].

Delta-band power for each channel was computed by first applying a band-pass filter within the 1-4Hz range and then calculating the amplitude of the Hilbert-transformed signal. Then the power for each channel is individually normalized by subtracting its mean and dividing by its standard deviation. The averaged delta power is obtained by taking the mean across all the recording channels.

#### 3.6 Local Field Potential Preprocessing

##### 3.6.1 Delta-band Power

Delta power were computed for local field potentials (LFPs) across all recorded channels. To calculate delta power, a band-pass filter (1-4 Hz) was applied to the LFP signal of each channel, followed by rectification and lowpass filtering ($<$ 0.72 Hz, corresponding to $\pi$ cycles of the mean band-pass frequencies).

##### 3.6.2 Hippocampal Sharp Wave Ripples (SWRs)

Hippocampal sharp wave ripples (SWRs) are brief, high-frequency oscillations (110-200Hz) that can be observed in the local field potential (LFP) recorded from hippocampal recording sites. For ripple detection in this study, we employed an offline method [16, 17] utilizing the LFP signal (1250 Hz) captured from the hippocampal CA1 region. The identification of ripple events was conducted individually for each CA1 recording site (channel), resulting in robust and extensively overlapping ripple detection across the channels. To consolidate the detection outcomes from various channels, a criterion was imposed: a detected ripple event was deemed valid only if it was identified in more than 40% of the CA1 channels.

#### 3.7 Semantic Decoding Model

The semantic decoding model is designed to evaluate the semantic information contained in the stimuli-evoked BOLD responses, generating text captions that describe the image stimuli presented to the subject. The decoding model consists of two main components: an fMRI encoder and a caption decoder (Figure 2A).

##### 3.7.1 fMRI Encoder

The fMRI encoder is used to extract latent representation from the BOLD response. The detailed architecture of the fMRI encoder, as shown in Figure S5A, utilizing convolutional layers and residual connections, aims to transform the high-dimensional BOLD response into 512-dimensional fMRI embeddings. To address the challenge of fMRI data scarcity, the fMRI encoder was trained to align the fMRI embeddings with the CLIP embedding space which has been extensively pre-trained using 400 million (image, text) pairs, thus offering a rich, 512-dimensional target. Therefore, we trained the fMRI encoder in contrastive learning paradigm [18, 19], aiming to maximize the alignment between the fMRI embeddings and the corresponding CLIP text embeddings, while minimizing the alignment with mismatched pairing. To achieve this, we use the contrastive training loss with loss function defined for $i^{\text{th}}$ fMRI embedding $Z_i$ and $j^{\text{th}}$ CLIP text embedding $T_j$ within a batch $B$ as:

$$\mathcal{L}_{contrast}(Z_i, T_j) = -\log \frac{\exp(\text{sim}(Z_i, T_j)/\tau)}{\sum_{i,j \in B, i \neq j} \exp(\text{sim}(Z_i, T_j)/\tau)} \tag{1}$$

Here, $\tau$ represents the temperature, a hyperparameter hyperparameter, and $\text{sim}(\cdot, \cdot)$ computes vector similarity, with cosine similarity being applied in this instance:

$$\mathcal{L}_{contrast}(Z_i, T_j) = -\log \frac{\exp(\cos(Z_i, T_j)/\tau)}{\sum_{i,j \in B, i \neq j} \exp(\cos(Z_i, T_j)/\tau)}. \tag{2}$$

In addition, we also maximize the alignment between fMRI embeddings and CLIP embeddings by incorporating the cosine loss defined as

$$\mathcal{L}_{align}(Z_i, T_i) = \lambda_1(1 - \cos(Z_i, T_i)) \tag{3}$$

Therefore, the total loss is defined as the summation of alignment loss and contrastive loss:

$$\mathcal{L}_{total} = \mathcal{L}_{align} + \lambda_2 \mathcal{L}_{contrast} \tag{4}$$

where $\lambda_1$ and $\lambda_2$ are tuning hyperparameters. In our training regimen, we set $\lambda_1 = 0.35$, $\lambda_2 = 0.65$ and $\tau = 0.45$. We also incorporate dropout before the final layers, with a dropout ratio of 0.3.

###### 3.7.2 Caption Decoder

We utilized a pre-trained caption decoder, DeCap [20], to generate captions from fMRI embeddings. DeCap was initially trained to produce captions using CLIP text embeddings based on large text corpus. Since the fMRI embeddings were aligned with the CLIP space through the fMRI encoder, we directly employed DeCap to decode these fMRI embeddings, for the generation of captions that describe the content of image stimuli shown to the subjects.

###### 3.7.3 Representation Similarity Analysis

We analyzed the alignment between CLIP text embeddings and fMRI embeddings through representation similarity analysis (RSA). For each validation fold, we constructed representation dissimilarity matrices (RDMs) for both the fMRI and CLIP text embeddings. To facilitate visualization and interpretation of the RDMs' structure, we employed t-SNE techniques [21] to project CLIP text embeddings into a 2-dimensional space. Subsequently, we applied k-means clustering to group stimuli with minimal Euclidean distance in this 2-dimensional representation into the same cluster. The RDMs for both fMRI and CLIP text embeddings, as depicted in Figure 2A, are organized and sorted according to this clustering scheme. The similarity between two RDMs is quantified using Pearson's correlation coefficient.

###### 3.7.4 Semantic Similarity Metrics

To evaluate the fidelity of generated captions to their semantic content, we computed semantic similarity score between the predicted captions and ground truth captions labeled by human. This evaluation employs several established metrics widely used in computer vision and natural language processing research: Bleu_1, Bleu_2 [22], CIDEr [23], METEOR [24], and ROUGE_L [25], with each of these metrics offering a different perspective on the semantic alignment between generated and ground truth text. Given the unique limitation inherent to each metric, we additionally defined a composite semantic metric (CSS) derived by averaging the scores from the aforementioned metrics, thereby providing a more holistic evaluation of caption fidelity.

To assess the statistical significance of the model-generated captions, for each trial/image, the semantic similarity score, for example, CSS score, is compared against a null distribution. This null distribution is constructed from the CSS scores between the ground truth caption and 1,000 randomly generated captions, which are generated by randomly sampling from the CLIP embedding space [18]. Notably, this null distribution exhibits variability across different trials, reflecting the diverse complexity levels associated with the semantic content of each image stimulus. Such approach ensures the evaluations accurately reflecting the model's performance in generating semantically coherent captions and invariant to the varying degrees of semantic complexity present across trials.

The same evaluative framework is applied across all aforementioned metrics to assess the accuracy of the generated captions. Encoding accuracy is quantified as the proportion of trials that are significant relative to null distributions, using a significance level of 0.95.

###### 3.7.5 Region-wise Decoding

In our study, unless otherwise specified, we use the decoding model to decode the response of voxels within the "nsdgeneral" ROI which includes occipital regions that are generally responsive in the NSD experiment [4].

For regionwise decoding analysis as shown in Figure 4 and 10, we adopted the regions of interest (ROIs) as delineated by the Yeo-17 network [13]. For each specific ROI, only the voxels that are defined by the corresponding ROI mask are considered. The responses from these selected voxels are then utilized as inputs to the decoding model to generate captions.

##### 3.8 Memory Encoding and Recall

The NSD dataset includes visual memory tasks, where each image stimulus was presented to participants three times. During each presentation, participants are prompted to indicate whether they have previously seen the image. By dissecting the memory task performance, we derived two memory measurements for evaluating the memory function based on participant response: memory encoding and memory recall.

**Memory encoding**, as a proxy for measuring the sensory coding efficacy, tends to evaluate how effectively the participant encodes a novel image into memory such that the participant can correctly recognize the image as previously seen. We formally define the memory encoding accuracy for each novel image as the memory recognition accuracy at the second presentation of the image.

**Memory recall**, on the other hand, tends to evaluate how effectively the participants is able to accurately retrieve and recognize an image previously seen. For each image, we can derive two recall accuracy, corresponding to the participant's recognition performance during the second and third presentations of the image, which is not novel to the participant.

##### 3.9 Neural Population Sensory Decoding Analysis

To assess the sensory information encoded by neuronal populations, we analyzed spiking data obtained during natural scene image stimulation sessions from the Allen Institute Visual Coding Mice Neuropixel dataset. In these sessions, mice were subjected to passive viewing of a sequence of images, each displayed for a duration of 250 milliseconds. Strictly following the approach in [26], we defined the neural code as a population response to each displayed image and employed support vector machines (SVMs) based on neural code to decode which one of the 118 images was viewed by the mouse. The sensory efficacy is quantified by the decoding accuracy of the SVMs.

##### 3.10 Infra-slow Propagating Waves

###### 3.10.1 Delay-pofile Decomposition

Following prior studies [17, 27] for analyzing the temporal dynamics of neural signals, we apply delay-profile decomposition method to identify the activation phase of particular regions or neuron that are activated in relation to specific events. Briefly, for each candidate event $k$, we derived a **delay profile** vector $d_k \in \mathbb{R}^N$ representing the activation phases of each region or neuron within that event:

$$\boldsymbol{d}_k = \left(t_{1k}\ t_{2k}\ \cdots\ t_{Nk},\right)^T \tag{5}$$

where $t_{i,k}$ represents the temporal centroid of the signal within the time segment $k$ for the $i$-th region/neuron and $N$ represents the total number of regions/neurons.

To represent the entire set of temporal relationships among the regions/neurons relative to each event segment analyzed, we combined the delay profiles for all $M$ segments into a delay profile matrix $\boldsymbol{D} \in \mathbb{R}^{M \times N}$:

$$\boldsymbol{D} = \begin{pmatrix} \boldsymbol{d}_1 & \boldsymbol{d}_2 & \cdots & \boldsymbol{d}_M \end{pmatrix} \tag{6}$$

$$= \begin{pmatrix} t_{11} & t_{12} & \cdots & t_{1N} \\ t_{21} & t_{22} & \cdots & t_{2N} \\ \vdots & \vdots & & \vdots \\ t_{M1} & t_{M2} & \cdots & t_{MN} \end{pmatrix} \tag{7}$$

Singular value decomposition (SVD) was then applied to the delay matrix $\boldsymbol{D}$ with the resulting principle component $\boldsymbol{u}^* \in \mathbb{R}^N$ defined as the **principal delay profile (PD)** representing the major sequential organization among the collective regions or neurons.

###### 3.10.2 fMRI SM-to-DMN Propagating Waves

To detect SM-to-DMN propagating waves, we adopt a template-matching approach [27]. The propagations, which typically involve a majority of cortical regions, are assumed entrained within the fMRI global signal fluctuations. Accordingly, we defined set of candidate events based on the low-frequency (<0.15Hz) components of the global signal, with the event boundaries defined by the adjacent troughs. This segmentation ensures that each candidate event contains a peak of global neural activity flanked by periods of relative quiescence.

For each candidate event $k$, we derive a delay profile as described previously. A candidate event $k$ is then considered as SM-to-DMN propagation if its delay profile closely matches the principal gradient profile (PG) $\boldsymbol{u}_{PG} \in \mathbb{R}^N$ [28]. The degree of similarity is quantitatively assessed using Pearson's correlation coefficient:

$$r_k = \text{PearsonCorr}(\boldsymbol{d}_k, \boldsymbol{u}_{PG}) \tag{8}$$

The significance thus can be evaluated from the correlation $r_k$ and we consider a candidate event as SM-to-DMN propagation if the corresponding p-value less than 0.001.

###### 3.10.3 Spiking Cascade

Processed neural spike data sorted with Kilosort2 pipeline [29] from the Allen Institute Visual Coding Mice Neuropixel dataset were utilized in this analysis. In line with prior studies [17, 26], we first computed neural spike rates using 200ms time bins and identified candidate infra-slow neural events by segmenting the spiking rate data based on the troughs of the filtered global mean spike rate (low-pass, 0.25Hz). We then, using the delay-profile decomposition method, extract delay profile for each candidate event and the principal delay profile representing the predominant sequential pattern within infra-slow brain activity. A candidate event $k$ was considered as a valid cascade event when the principal delay (PD) profile $\boldsymbol{u}^* \in \mathbb{R}^N$ matched with its delay profile $\boldsymbol{d}_k \in \mathbb{R}^N$, as quantified by Pearson's correlation coefficient:

$$r_k = \text{PearsonCorr}(\boldsymbol{d}_k, \boldsymbol{u}^*) \tag{9}$$

A candidate event that has a significant $r_k$ with $p < 0.001$ was considered a cascade event in our study.

Adhering to the analysis in previous studies [17, 26], we categorized neurons into two distinct groups based on their positioning within the PD profile: **positive-delay neurons** exhibiting significant PD values and **negative-delay neurons** characterized by significantly negative PD values ($p < 0.001$, one-sample t-test).

The slow cascade features a sharp increase in the spiking activity of positive-delay neurons in the middle, following the definition in [17], we thus identified and defined these time points as the local peak of the first-order temporal derivative of the mean spiking time course of the positive-delay neurons.

###### 3.10.4 Modulation Across Propagating Wave Cycles

To assess the fluctuation of the trial-based measurements, i.e., semantic encoding accuracy, memory encoding/decoding accuracy across the propagating waves cycle, as shown in Figure

3E, we constructed time series $f_{acc}(t)$ for each measurement during fMRI scan session defined as

$$f_{acc}(t) = \sum_{k=1}^{n} f_{acc}^{(k)}(t) \tag{10}$$

$$\text{with } f_{acc}^{(k)}(t) = \begin{cases} C_k & \text{if } T_k \leq t \leq T_k + \Delta T \\ NaN & \text{otherwise,} \end{cases} \tag{11}$$

where $T_k$ is the onset time of stimulus and $C_k$ is the accuracy for the k-th trial, and $\Delta T$ is the time window following the stimulus onset, which was set to 2 seconds throughout our study. The NaN stands for "Not a Number" and is omitted in the summation in equation 10.

Therefore, by formulating these trial-based assessments as time series, they are analyzed exactly same to other continuous measurements, such as pupil diameter and delta power. For the analysis of modulation across the propagation wave cycle, we aligned and averaged these time-series metrics relative to the propagation center (defined as the global signal peak) and normalized to the change in percentage relative to baseline, defined from -21s to -11s prior to the global signal peak.

### References

[1] Krauth, A., Blanc, R., Poveda, A., Jeanmonod, D., Morel, A., Székely, G.: A mean three-dimensional atlas of the human thalamus: generation from multiple histological data. Neuroimage **49**(3), 2053–2062 (2010)

[2] Edlow, B.L., Takahashi, E., Wu, O., Benner, T., Dai, G., Bu, L., Grant, P.E., Greer, D.M., Greenberg, S.M., Kinney, H.C., *et al.*: Neuroanatomic connectivity of the human ascending arousal system critical to consciousness and its disorders. Journal of Neuropathology & Experimental Neurology **71**(6), 531–546 (2012)

[3] Van Essen, D.C., Ugurbil, K., Auerbach, E., Barch, D., Behrens, T.E., Bucholz, R., Chang, A., Chen, L., Corbetta, M., Curtiss, S.W., *et al.*: The human connectome project: a data acquisition perspective. Neuroimage **62**(4), 2222–2231 (2012)

[4] Allen, E.J., St-Yves, G., Wu, Y., Breedlove, J.L., Prince, J.S., Dowdle, L.T., Nau, M., Caron, B., Pestilli, F., Charest, I., *et al.*: A massive 7t fmri dataset to bridge cognitive neuroscience and artificial intelligence. Nature neuroscience **25**(1), 116–126 (2022)

[5] Gu, Y., Han, F., Sainburg, L.E., Schade, M.M., Buxton, O.M., Duyn, J.H., Liu, X.: An orderly sequence of autonomic and neural events at transient arousal changes. Neuroimage **264**, 119720 (2022)

[6] Siegle, J.H., Jia, X., Durand, S., Gale, S., Bennett, C., Graddis, N., Heller, G., Ramirez, T.K., Choi, H., Luviano, J.A., *et al.*: Survey of spiking in the mouse visual system reveals functional hierarchy. Nature **592**(7852), 86–92 (2021)

[7] Vries, S.E., Lecoq, J.A., Buice, M.A., Groblewski, P.A., Ocker, G.K., Oliver, M., Feng, D., Cain, N., Ledochowitsch, P., Millman, D., *et al.*: A large-scale standardized physiological survey reveals functional organization of the mouse visual cortex. Nature neuroscience **23**(1), 138–151 (2020)

[8] Steinmetz, N.A., Zatka-Haas, P., Carandini, M., Harris, K.D.: Distributed coding of choice, action and engagement across the mouse brain. Nature **576**(7786), 266–273 (2019)

[9] Glasser, M.F., Sotiropoulos, S.N., Wilson, J.A., Coalson, T.S., Fischl, B., Andersson, J.L., Xu, J., Jbabdi, S., Webster, M., Polimeni, J.R., *et al.*: The minimal preprocessing pipelines for the human connectome project. Neuroimage **80**, 105–124 (2013)

[10] Salimi-Khorshidi, G., Douaud, G., Beckmann, C.F., Glasser, M.F., Griffanti, L., Smith, S.M.: Automatic denoising of functional mri data: combining independent component analysis and hierarchical fusion of classifiers. Neuroimage **90**, 449–468 (2014)

[11] Kay, K., Rokem, A., Winawer, J., Dougherty, R., Wandell, B.: Glmdenoise: a fast, automated technique for denoising task-based fmri data. Frontiers in neuroscience, 247 (2013)

[12] Biswal, B.B., Mennes, M., Zuo, X.-N., Gohel, S., Kelly, C., Smith, S.M., Beckmann, C.F., Adelstein, J.S., Buckner, R.L., Colcombe, S., *et al.*: Toward discovery science of human brain function. Proceedings of the national academy of sciences **107**(10), 4734–4739 (2010)

[13] Schaefer, A., Kong, R., Gordon, E.M., Laumann, T.O., Zuo, X.-N., Holmes, A.J., Eickhoff, S.B., Yeo, B.T.: Local-global parcellation of the human cerebral cortex from intrinsic functional connectivity mri. Cerebral cortex **28**(9), 3095–3114 (2018)

[14] Liu, Z., Zwart, J.A., Gelderen, P., Kuo, L.-W., Duyn, J.H.: Statistical feature extraction for artifact removal from concurrent fmri-eeg recordings. Neuroimage **59**(3), 2073–2087 (2012)

[15] Falahpour, M., Chang, C., Wong, C.W., Liu, T.T.: Template-based prediction of vigilance

fluctuations in resting-state fmri. Neuroimage **174**, 317–327 (2018)

[16] Stark, E., Roux, L., Eichler, R., Senzai, Y., Royer, S., Buzsáki, G.: Pyramidal cell-interneuron interactions underlie hippocampal ripple oscillations. Neuron **83**(2), 467–480 (2014)

[17] Liu, X., Leopold, D.A., Yang, Y.: Single-neuron firing cascades underlie global spontaneous brain events. Proceedings of the National Academy of Sciences **118**(47), 2105395118 (2021)

[18] Radford, A., Kim, J.W., Hallacy, C., Ramesh, A., Goh, G., Agarwal, S., Sastry, G., Askell, A., Mishkin, P., Clark, J., *et al.*: Learning transferable visual models from natural language supervision. In: International Conference on Machine Learning, pp. 8748–8763 (2021). PMLR

[19] Lin, S., Sprague, T., Singh, A.K.: Mind reader: Reconstructing complex images from brain activities. Advances in Neural Information Processing Systems **35**, 29624–29636 (2022)

[20] Li, W., Zhu, L., Wen, L., Yang, Y.: Decap: Decoding clip latents for zero-shot captioning via text-only training. arXiv preprint arXiv:2303.03032 (2023)

[21] Maaten, L., Hinton, G.: Visualizing data using t-sne. Journal of machine learning research **9**(11) (2008)

[22] Papineni, K., Roukos, S., Ward, T., Zhu, W.-J.: Bleu: a method for automatic evaluation of machine translation. In: Proceedings of the 40th Annual Meeting of the Association for Computational Linguistics, pp. 311–318 (2002)

[23] Vedantam, R., Lawrence Zitnick, C., Parikh, D.: Cider: Consensus-based image description evaluation. In: Proceedings of the IEEE Conference on Computer Vision and Pattern Recognition, pp. 4566–4575 (2015)

[24] Denkowski, M., Lavie, A.: Meteor universal: Language specific translation evaluation for any target language. In: Proceedings of the Ninth Workshop on Statistical Machine Translation, pp. 376–380 (2014)

[25] Lin, C.-Y.: Rouge: A package for automatic evaluation of summaries. In: Text Summarization Branches Out, pp. 74–81 (2004)

[26] Yang, Y., Leopold, D.A., Duyn, J., Sipe, G., Liu, X.: Intrinsic forebrain arousal dynamics governs sensory stimulus encoding. bioRxiv, 2023–10 (2023)

[27] Gu, Y., Sainburg, L.E., Kuang, S., Han, F., Williams, J.W., Liu, Y., Zhang, N., Zhang, X., Leopold, D.A., Liu, X.: Brain activity fluctuations propagate as waves traversing the cortical hierarchy. Cerebral cortex **31**(9), 3986–4005 (2021)

[28] Margulies, D.S., Ghosh, S.S., Goulas, A., Falkiewicz, M., Huntenburg, J.M., Langs, G., Bezgin, G., Eickhoff, S.B., Castellanos, F.X., Petrides, M., *et al.*: Situating the default-mode network along a principal gradient of macroscale cortical organization. Proceedings of the National Academy of Sciences **113**(44), 12574–12579 (2016)

[29] Stringer, C., Pachitariu, M., Steinmetz, N., Reddy, C.B., Carandini, M., Harris, K.D.: Spontaneous behaviors drive multidimensional, brainwide activity. Science **364**(6437), 7893 (2019)
